# Harnessing Lytic Phages for Biofilm Control in Carbapenem-Resistant *Klebsiella pneumoniae* Causing Urinary Tract Infection

**DOI:** 10.1101/2025.10.01.679830

**Authors:** Dipendra Kumar Mandal, Elisha Upadhaya, Puja Dahal, Gaurav Adhikari, Rojina Pandey, Abdul Rehaman, Sudip Timilsina, Pragya Sapkota, Shobha Amagain, David Pun, Kundan Khadka, Keshab Gorathoki, Bijita Neupane, Sangharsika Chaudhary, Sushila Thapa, Binod Khadka, Gun Raj Dhungana, Gorkha Raj Giri, Roshan Nepal, Rajindra Napit, Pragati Pradhan, Krishna Das Manandhar, Rajani Malla

## Abstract

**Background:** *Klebsiella pneumoniae* is a major opportunistic pathogen with rising multidrug resistance and biofilm-related infections. Molecular and phage characterization is crucial to understand resistance mechanisms and explore alternative therapies such as phage therapy.

**Methods:** We performed whole-genome sequencing and antibiotic susceptibility testing of hospital-isolated *Klebsiella pneumoniae* (KP6697). MLST, plasmid replicon analysis, and resistance gene identification were conducted using bioinformatics. Phage isolation, electron microscopy-based morphological and biofilm analysis, and evaluation of lytic activity, stability, and host range were performed. Phage genome sequencing and annotation identified functional genes.

**Results:** The host strain *Klebsiella pneumoniae* (KP6697) was multidrug-resistant, exhibiting resistance to 18 of 22 tested antibiotics, and genome analysis identified ST16 with eight plasmid replicons and 23 resistance genes, including blaCTX-M-15, blaNDM-5, and blaOXA-181. Functional annotations revealed extensive metabolic versatility and a rich repertoire of genes for biofilm formation, quorum sensing, secretion systems, and stress response. A lytic phage, Phage_KP6697_Omshanti, was isolated and classified as a Caudoviricetes member with a 45.3kb genome encoding lysis, replication, and structural genes. It demonstrated short latency, high burst size, thermal and pH stability, and broad host range against CRKP and other MDR strains. Importantly, microscopy confirmed its ability to inhibit and degrade biofilms at multiple stages, highlighting strong therapeutic potential.

**Conclusion:** Comprehensive analysis of carbapenem-resistant *K. pneumoniae* (KP6697) revealed multidrug resistance and strong biofilm formation. The lytic phage Phage_KP6697_Omshanti, with depolymerase and endolysin activity, disrupted biofilms, and its stability, high burst size, and genomic traits suggest potential as an anti-CRKP agent, especially with antibiotics

**IMPORTANCE:** *Klebsiella pneumoniae* is increasing multidrug resistance and robust biofilm formation pose severe clinical challenges, limiting treatment options. Understanding the molecular basis of its resistance and exploiting bacteriophages with strong biofilm-disrupting properties provide promising alternative therapeutic strategies. This study highlights the isolation and genomic characterization of a lytic phage with potent anti-biofilm activity against carbapenem-resistant *K. pneumoniae*, underscoring its potential in combating resistant infections.

## Introduction

*Klebsiella pneumoniae* is one of the six challenging pathogens grouped under the ESKAPE (*Enterococcus faecium*, *Staphylococcus aureus*, *Klebsiella pneumoniae*, *Acinetobacter baumannii*, *Pseudomonas aeruginosa*, and *Enterobacter* species) acronym due to their multidrug resistance and clinical relevance (1). Carbapenem-resistant *Klebsiella pneumoniae* (CRKP) has emerged globally as a critical threat largely driven by plasmid mediated carbapenemase genes that facilitate horizontal transfer across bacterial population. Alarmingly, CRKP strains have begun showing resistance to last-resort antibiotics like polymyxin and tigecycline (2). Although new β-lactam/β-lactamase inhibitor combinations such as cefatazidime/avibactam, meropenem/vaborbactam and imipenem/relebactam have been introduced, therapeutic options remain limited, promoting exploration of alternative strategies such as bacteriophage therapy.

A critical aspect of CRKP pathogenicity lies in its ability to form robust biofilms on both biotic and abiotic surfaces. Biofilms are structured microbial communities encased in a self-produced extracellular polymeric matrix, providing bacteria with enhanced survival advantages in hostile environments (3, 4). Within healthcare settings, CRKP biofilms colonize medical devices including urinary catheters, central venous lines, endotracheal tubes, and prosthetic implants, creating persistent sources of infection that are notoriously difficult to eradicate (5–7).

The biofilm lifestyle confers multiple advantages to CRKP, including protection from host immune responses, reduced susceptibility to antimicrobial agents, and enhanced horizontal gene transfer capabilities (8, 9). Biofilm-associated bacteria exhibit tolerance to antibiotic concentrations up to 1000-fold higher than their planktonic counterparts through various mechanisms including limited drug penetration, altered metabolic states, and the presence of persister cells (10, 11). This phenotypic resistance, combined with genetically encoded resistance mechanisms, creates a formidable barrier to successful treatment of CRKP infections. The molecular machinery underlying biofilm formation in CRKP involves complex regulatory networks controlling adhesion, matrix synthesis, and community development. Key components include polysaccharide synthesis pathways (such as poly-β-1,6-N-acetyl-D-glucosamine production), fimbrial adhesion systems (including type 1 and type 3 fimbriae), and sophisticated two-component regulatory systems that respond to environmental stimuli (3, 12–14). Understanding these biofilm formation mechanisms is crucial for developing targeted therapeutic interventions that can disrupt established biofilm communities.

Bacteriophages (or phages) are viruses that specifically infect bacteria and particularly lytic phages can lyse bacterial cells. Because of its bactericidal property phages are considered as an alternative measures to prevent and control various bacterial infections including CRKP infections (15, 16). Phage therapy, predating antibiotics by nearly a century, fell out of favor with antibiotic proliferation but has recently regained attention within rising antibiotic resistance (17–19). Clinically, phages can be administered systemically or locally and have shown promise against *Enterobacteriaceae* pathogens, with *Escherichia* phage and *Salmonella* phages being well studied (20–22) Applications of phages against *K. pneumoniae* have been documented in burn wounds and diabetic foot ulcers (23, 24). However, data on phage efficacy against clinical CRKP strains remain scarce.

The rising resistance of CRKP, the role of biofilms in persistent infections, and the potential of bacteriophages highlight the need for innovative treatments. This study bridges the gap between CRKP biofilm formation and phage-mediated disruption by analyzing the molecular mechanisms of biofilm development and characterizing a novel lytic bacteriophage, Phage_KP6697_Omshanti, effective against CRKP biofilms. Functional genomic analysis revealed 87 genes involved in biofilm formation, covering structural, regulatory, and metabolic pathways. The phage demonstrated both anti-CRKP activity and biofilm disruption capabilities. This dual approach elucidating biofilm mechanisms while developing targeted phage therapeutics offers a new strategy for combating antibiotic-resistant infections. Identifying biofilm formation pathways reveals therapeutic targets, while characterizing anti-biofilm phages provides immediate clinical applications and informs the design of enhanced phage therapies. This is critical for CRKP infections, often involving treatment-resistant biofilms. The narrow host range of the phage addresses concerns about off-target effects, enhancing treatment efficacy. By combining mechanistic insights with practical therapeutic development, this research provides a foundation for targeted interventions, phage cocktails, and engineered therapies to tackle CRKP biofilm infections, addressing a major challenge in infectious disease medicine.

## Results

### Antibiotic resistance genes profile of the host bacteria

Antibiotic susceptibility of host bacteria *Klebsiella pneumoniae* (KP6697) was determined by Kirby-Bauer disk diffusion method and is elaborated in (Table 1). The isolate revealed resistance to 18 out of the 22 antibiotics tested. Based on resistance to at least one agent in three or more antibiotic categories (25) and thus KP6697 was considered as a multidrug-resistant (MDR) isolate.

**Table 1.** Antibiotic susceptibility test by disc diffusion method of *K. pneumoniae* (KP6697). Interpretations are made according to the CLSI breakpoint tables.

| SN | Antibiotics | Code | MIC<br>(Vitek<br>II) | Inhibition<br>zone (in<br>mm) | Interpretation |
| --- | --- | --- | --- | --- | --- |
| 1 | Chloramphenicol (30µg/ml) | C | ≥8 | 10 | Resistant |
| 2 | Amoxycillin (10µg/ml) | AMP | ≥32 | 0 | Resistant |
| 3 | Tigecycline (15µg/ml) | TGC | ≥2 | 13 | Sensitive |
| 4 | Doxycycline (30µg/ml) | DO | ≥2 | 14 | Resistant |
| 5 | Ciprofloxacin (5µg/ml) | CIP | ≥4 | 0 | Resistant |
| 6 | Gentamicin (10µg/ml) | GEN | ≥16 | 0 | Resistant |
| 7 | Cotrimoxazole (14µg/ml) | COT | ≥320 | 0 | Resistant |
| 8 | Ceftazidime (14µg/ml) | CAZ | ≥64 | 0 | Resistant |
| 9 | Meropenem (10µg/ml) | MEM | ≥16 | 0 | Resistant |
| 10 | Piperacillin tazobactam | PIT | ≥128 | 0 | Resistant |
|  | (100/10µg/ml) |  |  |  |  |
| 11 | Nitrofurantoin (300µg/ml) | NIT | 128 | 13 | Resistant |
| 12 | Cefepime (30µg/ml) | CPM | ≥64 | 0 | Resistant |
| 13 | Cefoperazone sulbactam<br>(75/30µg/ml) | CPZ | ≥64 | 0 | Resistant |
| 14 | Imipenem (10µg/ml) | IMP | ≥16 | 0 | Resistant |
| 15 | Nalidixic acid (30µg/ml) | NA | ≥32 | 0 | Resistant |
| 16 | Aztreonam (30µg/ml) | AT | ≥64 | 8 | Resistant |
| 17 | Levofloxacin (5µg/ml) | LE | ≥8 | 10 | Resistant |
| 18 | Colistin (10µg/ml) | CL | <0.5 | - | Sensitive |
| 19 | Fosfomycin (200µg/ml) | FO | <16 | 18 | Sensitive |
| 20 | Amikacin (30µg/ml) | AK | ≤2 | 19 | Sensitive |
| 21 | Ceftriaxone (30µg/ml) | CTR | ≥64 | 17 | Resistant |
| 22 | Amoxy-clavulanic acid<br>(20/10µg/ml) | AMC | ≥32 | 12 | Resistant |

### Detection of Metallo-β-lactamase (MBL) production

The KP6697 isolate was confirmed MBL-positive by disk diffusion and combination disk methods. NDM-mediated carbapenem resistance was verified by Bio-Rad CFX96 PCR System targeting the *bla*NDM gene (∼621 bp) using in-house primers: NDM For (5’ CGGAATGGCTCATCACGATC-3’) and NDM Rev (5’ GGTTTGGCGATCTGGTTTTC-3’).

PCR conditions were: 3 min 30 sec at 95°C; 34 cycles of 30 sec at 60.4°C and 1 min at 72°C; followed by 5 min at 72°C, with the entire process completed in under 2hr. (Fig S1).

### Molecular characterization of host *K. pneumoniae* (KP6697)

The whole genome sequencing revealed that the isolate had the genome size of ∼4.9Mbp and based on the MLST typing, the isolate belonged to Sequence Type 16 (ST16) *Klebsiella*. Plasmid prediction identified a total of 8 replicons in this isolate. The plasmids present in the isolates were Col440II, ColKP3, IncFIA(HI1), IncFIB(K), IncFII, IncFII, IncR, and IncX3 (Table 2). The analysis performed for the presence of genes for antibiotic resistance revealed the presence of 23 different antibiotic resistance genes (Table 2). The major groups of antibiotics against which the resistance genes were present were beta-lactams (ampicillin), phenicols (chloramphenicol), aminoglycosides (gentamicin, amikacin and streptomycin), sulfonamides (trimethoprim and sulfamethoxazole), macrolides (erythromycin), fluoroquinolones (ciprofloxacin) and rifamycin (rifampicin). The genes involved in antibiotic resistance were *blaCTX-M-15, blaNDM-5, blaOXA-181, blaSHV-61, blaTEM-214* (beta-lactams), *catA1, cmlA1* (phenicols), *aac* (*3*)*-Iid, aadA2, ant (3’’)-Ia, rmtB* (aminoglycosides), *ere(A), erm(B), mph(A)* (macrolides), *dfrA12, sul1* (sulfonamides), *qnrS1* (fluoroquinolones), *ARR-3* (rifampicin) and *OqxA, OqxA, OqxB, OqxB, qacE* (Table 2). Furthermore, the CkeckV analysis detected 2 prophage sequences with ∼100% complete genomes of predicted genome size 39 and 40 Kbp. The prophage sequences were not subjected to any further analysis in this research.

**Table 2.**
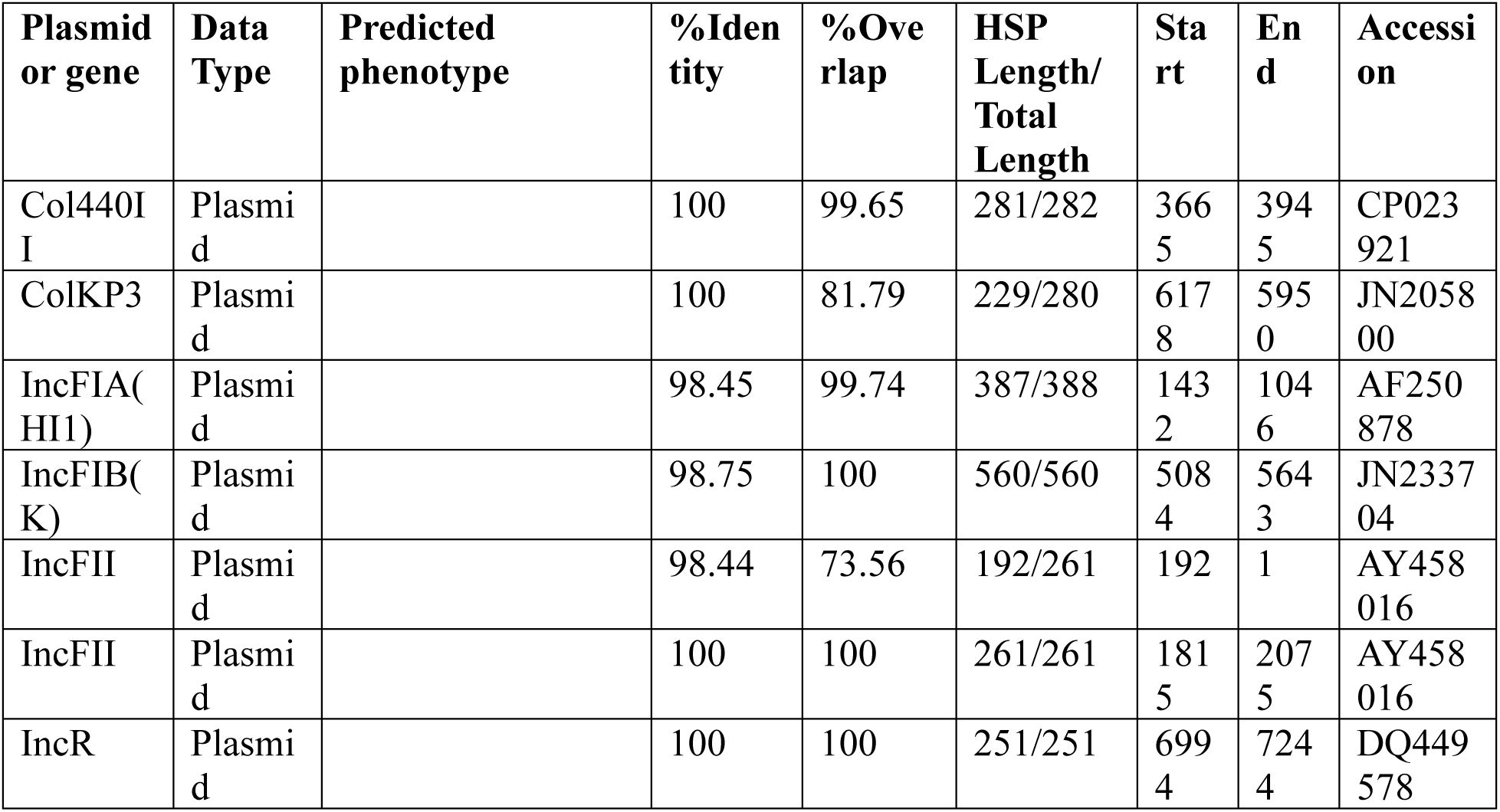
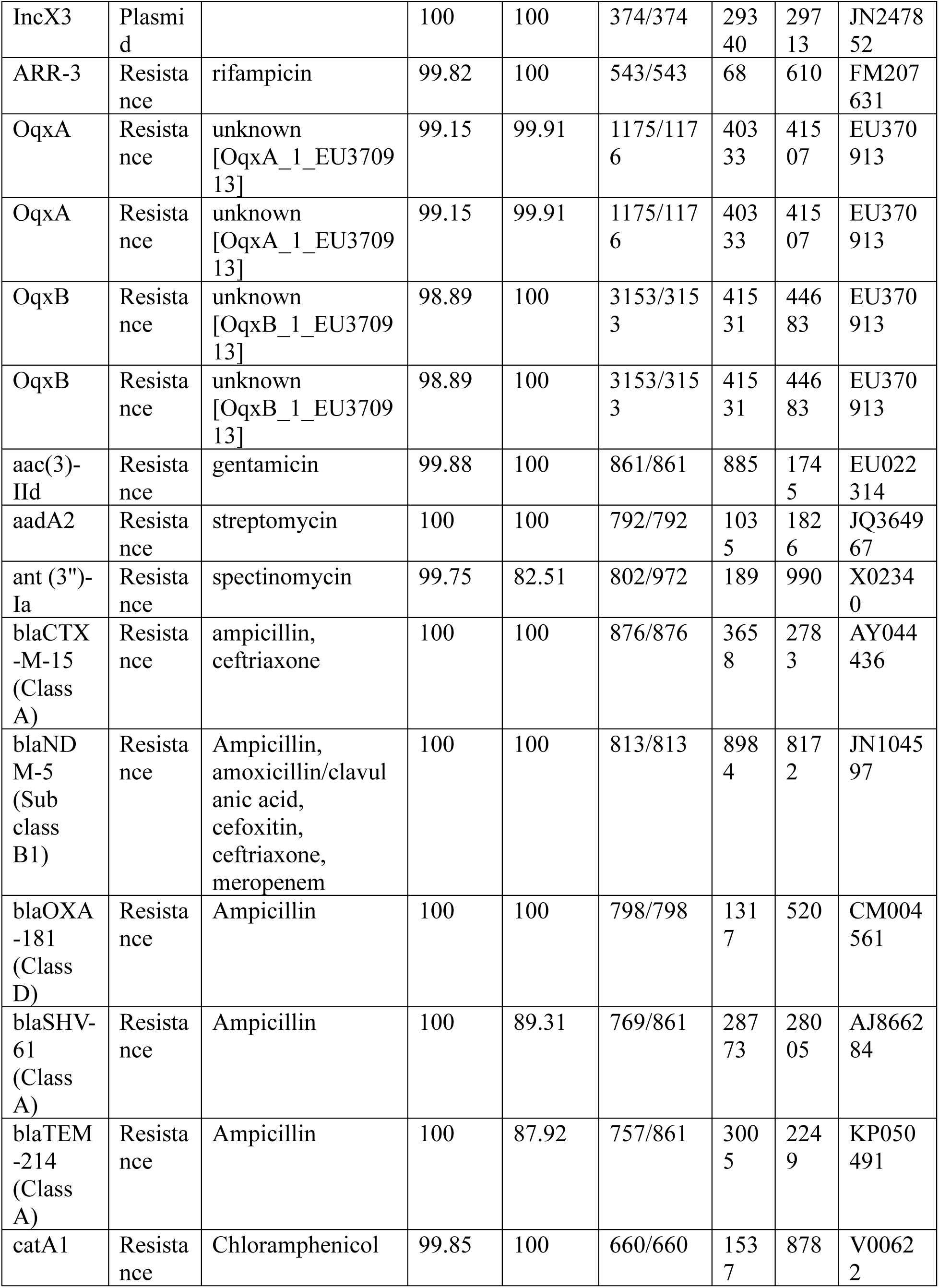
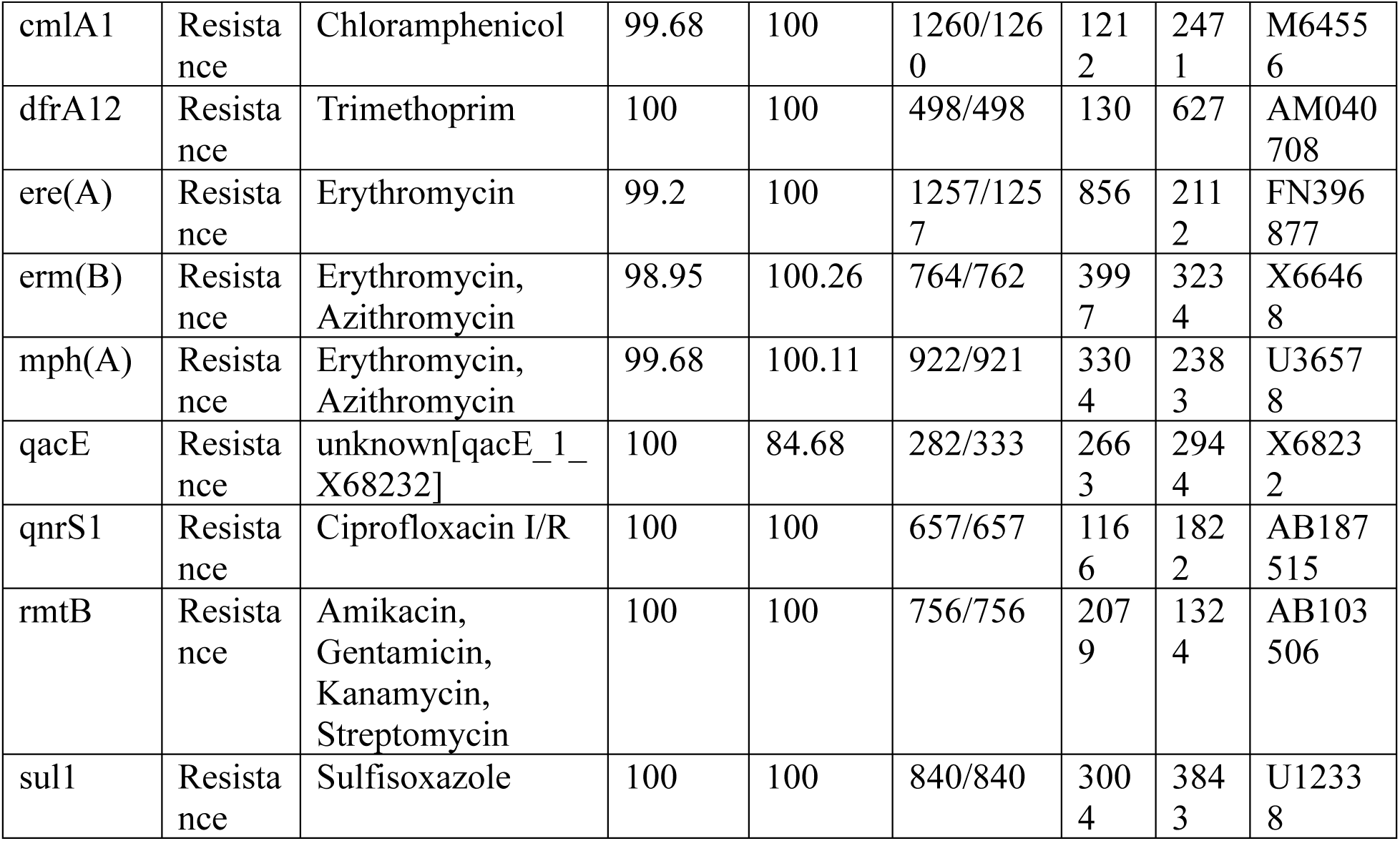
Molecular characteristics of *Klebsiella pneumoniae* KP6697 based on RESFINDER software.

### Functional annotations and characteristics of bacterial host KP6697

The genomic analysis of KP6697 with KEGG (Kyoto Encyclopedia of Genes and Genomes) database, a KO number (or K number) revealed a comprehensive set of metabolic and resistance pathways, as detailed in the pathway modules (Table S1). The genome encoded complete pathways for central carbohydrate metabolism, including glycolysis (M00001, 9/9), gluconeogenesis (M00003, 7/7), the citrate cycle (M00009, 8/8), and the pentose phosphate pathway (M00004, 6/6). Additional carbohydrate degradation pathways, such as D-galacturonate (M00631, 5/5) and D-glucuronate (M00061, 5/5), and biosynthesis of glycogen (M00854, 2/2) and trehalose (M00565, 6/6) were fully represented. Energy metabolism included complete carbon fixation pathways (e.g., reductive citrate cycle, M00173, 10/10) and ATP synthesis modules (e.g., F-type ATPase, M00157, 1/1). Nitrogen and sulfur metabolism were supported by assimilatory and dissimilatory nitrate reduction (M00531, M00530, 2/2 each) and sulfate reduction (M00176, 2/2).

Lipid metabolism encompassed fatty acid biosynthesis (M00082, 2/2; M00083, 1/1) and degradation (M00087, 3/3), alongside phospholipid biosynthesis (M00093, 3/3). Nucleotide metabolism included complete de novo purine (M00048, 8/8) and pyrimidine (M00051, 3/3) biosynthesis. Amino acid metabolism covered biosynthesis and degradation of serine, threonine, cysteine, methionine, branched-chain amino acids, lysine, arginine, proline, and histidine, with all pathways complete (e.g., lysine biosynthesis, M00016, 9/9; histidine biosynthesis, M00026, 6/6). Glycan metabolism included lipopolysaccharide (M00060, 9/9) and cofactor/vitamin biosynthesis pathways, such as thiamine (M00127, 7/7), riboflavin (M00125, 7/7), and heme (M00121, 10/10). Terpenoid and polyketide biosynthesis pathways were fully represented, including C5 isoprenoid (M00096, 8/8) and type II polyketide (M00778, 3/3) biosynthesis. Xenobiotic degradation pathways, such as toluene (M00538, 3/3) and benzoate (M00551, 2/2), were also complete. The genome encoded robust drug resistance mechanisms, including carbapenem (M00851, 1/1), methicillin (M00625, 3/3), and multiple multidrug efflux pumps (e.g., MexAB-OprM, M00718, 2/2). Symbiosis (M00664, 4/4) and metabolic capacities like nitrate (M00615, 2/2) and sulfate-sulfur assimilation (M00616, 2/2) were fully present, highlighting the bacterium’s metabolic versatility, environmental adaptability, and resistance to antibiotics.

#### A. Extracellular transport and secretion systems

Several genes were identified as part of extracellular transport and secretion systems, critical for bacterial interaction with their environment. Notably, genes encoding components of the general secretion pathway (GSP) were prevalent, including K02451–K02464 (general secretion pathway proteins B–O), which facilitate the export of proteins across the bacterial membrane. For instance, K02454 (general secretion pathway protein E, encodes an ATP-dependent component [EC:7.4.2.8], essential for type II secretion system functionality. Additionally, K02504 (*hofB*) and K02505 (*hofC*) encode protein transport proteins, while K02507 (*hofQ*) supports outer membrane protein transport, collectively enabling the secretion of extracellular enzymes and toxins. The presence of K11935 (*pgaA*) and K11937 (*pgaD*), involved in biofilm poly-beta-1,6-N-acetyl-D-glucosamine (PGA) synthesis, underscores the bacterium’s capacity to form robust extracellular matrices, enhancing community stability and resistance.

#### B. Regulatory mechanisms in host bacteria KP6697

The analysis identified multiple regulatory genes, particularly those encoding two-component systems and transcriptional regulators which orchestrate social behaviors such as quorum sensing and environmental response. Genes such as K07659 (*ompR*), K07657 (*phoB*), K07662 (*cpxR*), K07684 (*narL*), K07712 (*glnG*), K07713 (*hydG*), and K07715 (*glrR*) encode response regulators of two-component systems from the OmpR and NtrC families, modulating responses to environmental cues like phosphate availability, nitrogen levels, and osmotic stress (Table S2) (26), (27). Additionally, luxR family transcriptional regulators (K04333, K07782) were detected, with K07782 (sdiA) linked to quorum-sensing regulation, suggesting a sophisticated signaling network for coordinating group behaviors. The presence of sigma factors (K03086, K03087, K03092) further indicates fine-tuned regulation of gene expression, with K03086 encoding the primary RNA polymerase sigma factor and K03092 encoding the sigma-54 factor, critical for stress response and nitrogen assimilation.

#### C. Social genes for quorum sensing and biofilm production

The genomic analysis of the host bacteria KP6697 revealed a diverse repertoire of social genes associated with extracellular functions, regulatory mechanisms and biofilm formation. The identified genes, annotated with their respective KEGG Orthology (KO) definitions and functional roles, are summarized below, highlighting their contributions to bacterial social behavior, environmental adaptation and pathogenicity (Table S2).

##### Biofilm-associated gene functions

The functional analysis of the identified KO genes revealed a comprehensive molecular machinery involved in biofilm formation and maintenance. There were total of 87 unique KO entries detected in the isolate representing diverse functional categories critical for biofilm development and regulation.

**i. Direct biofilm formation genes:** Several genes directly involved in biofilm synthesis were identified:

- **Poly-β-1,6-N-acetyl-D-glucosamine (PNAG) synthesis pathway**: K11935 (biofilm PGA synthesis protein PgaA), K11931 and K21478 (poly-beta-1,6-N-acetyl-D-glucosamine N-deacetylase), and K11937 (biofilm PGA synthesis protein PgaD) were present, indicating active PNAG biosynthesis machinery.
- **Colanic acid production**: K13650 (MqsR-controlled colanic acid and biofilm protein A) was identified, suggesting involvement in extracellular polysaccharide matrix formation.
**ii. Fimbrial and adhesion systems:** A significant enrichment of fimbrial genes was observed: **Type 1 fimbriae**: K07345 (major type 1 subunit fimbrin/pilin) appeared 6 times in the dataset, indicating high abundance of type 1 fimbrial components. **Minor fimbrial components**: K07348 (minor fimbrial subunit) and K07350 (minor fimbrial subunit) were present, supporting complete fimbrial assembly. **Specialized adhesins**: K21967 (Mat/Ecp fimbriae adhesin) was identified, representing specialized attachment mechanisms.
**iii. Regulatory networks:** Multiple transcriptional regulatory systems controlling biofilm formation were detected: **Quorum sensing regulation**: K07782 (LuxR family transcriptional regulator, quorum-sensing system regulator SdiA) and K04333 (LuxR family transcriptional regulator, csgAB operon transcriptional regulatory protein) were present. **Two-component systems**: Several two-component regulatory systems were identified including K07659 (OmpR), K07657 (PhoB), K07662 (CpxR), K07684 (NarL), K07712 (GlnG), K07713 (HydG), and K07715 (GlrR).
**iv. Secretion systems:** Critical protein secretion machinery was represented: **General secretion pathway (Type II)**: Complete Gsp machinery from K02451 (GspB) through K02465 (GspS) was identified, enabling secretion of biofilm-associated proteins. **Type VI secretion system**: K11907 (VasG protein) was present, indicating potential involvement in inter-bacterial competition within biofilms.
**v. Iron acquisition systems:** Multiple iron-siderophore transport components were detected: **Siderophore transport**: K23186, K23187, K23188, and K25111 representing permease and ATP-binding proteins for iron-siderophore uptake. **Siderophore biosynthesis**: K01252 (bifunctional isochorismate lyase/aryl carrier protein) and K08225 (enterobactin exporter) were identified. **Siderophore reception**: K16090 (catecholate siderophore receptor) and K10829 (ferric hydroxamate transport ATP-binding protein) were present.

Functional category distribution comprises of genes for Biofilm matrix synthesis (8.4% of total KOs), Adhesion and fimbriae (12.6% of total KOs), Transcriptional regulation (16.8% of total KOs), Iron homeostasis (14.7% of total KOs), Protein secretion (18.9% of total KOs), Cell envelope modification (11.6% of total KOs), Stress response (17.0% of total KOs).

#### D. Stress response and enzymatic functions

Genes encoding stress response and enzymatic functions were also identified, contributing to bacterial resilience and metabolic versatility. For instance, K04564 (superoxide dismutase, Fe-Mn family [EC:1.15.1.1]) protects against oxidative stress, while K18699 (beta-lactamase class A SHV [EC:3.5.2.6]) confers resistance to beta-lactam antibiotics. Proteases such as K04772 (DegQ [EC:3.4.21.-]) and K18988 (serine-type D-Ala-D-Ala carboxypeptidase/endopeptidase [EC:3.4.16.4 3.4.21.-]) likely contribute to cell wall remodeling and pathogenicity. Additionally, K07173 (S-ribosylhomocysteine lyase [EC:4.4.1.21]) and K06998 (trans-2,3-dihydro-3-hydroxyanthranilate isomerase [EC:5.3.3.17]) indicate metabolic pathways for quorum-sensing autoinducer synthesis and aromatic compound degradation respectively.

#### E. Other functional genes

Further analysis revealed additional genes with diverse roles, including K03590 (*ftsA*), involved in cell division, and K03806 (N-acetyl-anhydromuramoyl-L-alanine amidase [EC:3.5.1.28]), contributing to peptidoglycan turnover. The type VI secretion system protein K11907 (*vasG*) suggests potential for interbacterial competition or host interaction. Genes like K01649 (2-isopropylmalate synthase [EC:2.3.3.13]) and K00849 (galactokinase [EC:2.7.1.6]) indicate metabolic capabilities for amino acid and carbohydrate utilization respectively. In summary, the identified social genes highlight the bacterium’s ability to engage in complex extracellular interactions, regulate gene expression in response to environmental stimuli, form biofilms, acquire essential nutrients like iron and withstand stress. These genetic features collectively underscore the bacterium’s adaptive and social capabilities, likely contributing to its ecological success and potential pathogenicity.

### Harnessing lytic phage to control biofilm forming isolate KP6697

One *Klebsiella* Phage_KP6697_Omshanti against the bacterial host KP6697 was isolated and purified. The Phage_KP6697_Omshanti formed opaque halo zone around clear plaques and exhibited a potent lytic activity against the host KP6697. Phage_KP6697_Omshanti formed bull’s eye shaped clear plaques (3 mm in diameter) and initial number was three that were surrounded by large opaque halo zone (Fig S2).

Furthermore, the transmission electron microscopy revealed that the Phage_KP6697_Omshanti had an isometric, polyhedral head (88.6 nm in diameter) and a short tail (120 nm long) (Fig 1).

**Fig 1:** Phage_KP6697_Omshanti morphology under TEM

According to the current ICTV classification system, phage was classified as belonging to the Caudoviricetes class of the order Caudovirales.

### Growth and physical characteristics of Phage_KP6697

#### a) Optimal MOI and one-step growth curve

Based on the maximum phage titer obtained upon infection of host KP6697, the optimal MOI of Phage_KP6697_Omshanti was determined to be 0.1. The one-step growth curve of phage showed a latent period was 20 min and a burst period of 30 min (Fig 2), followed by a plateau period. The size of the burst, calculated (28) as the ratio of the average number of phages released per infected host cell before (2.66 ×10^8^ CFU) and after the burst (3.17×10^10^PFU), was approximately 119 PFU/cell.

**Fig 2:**
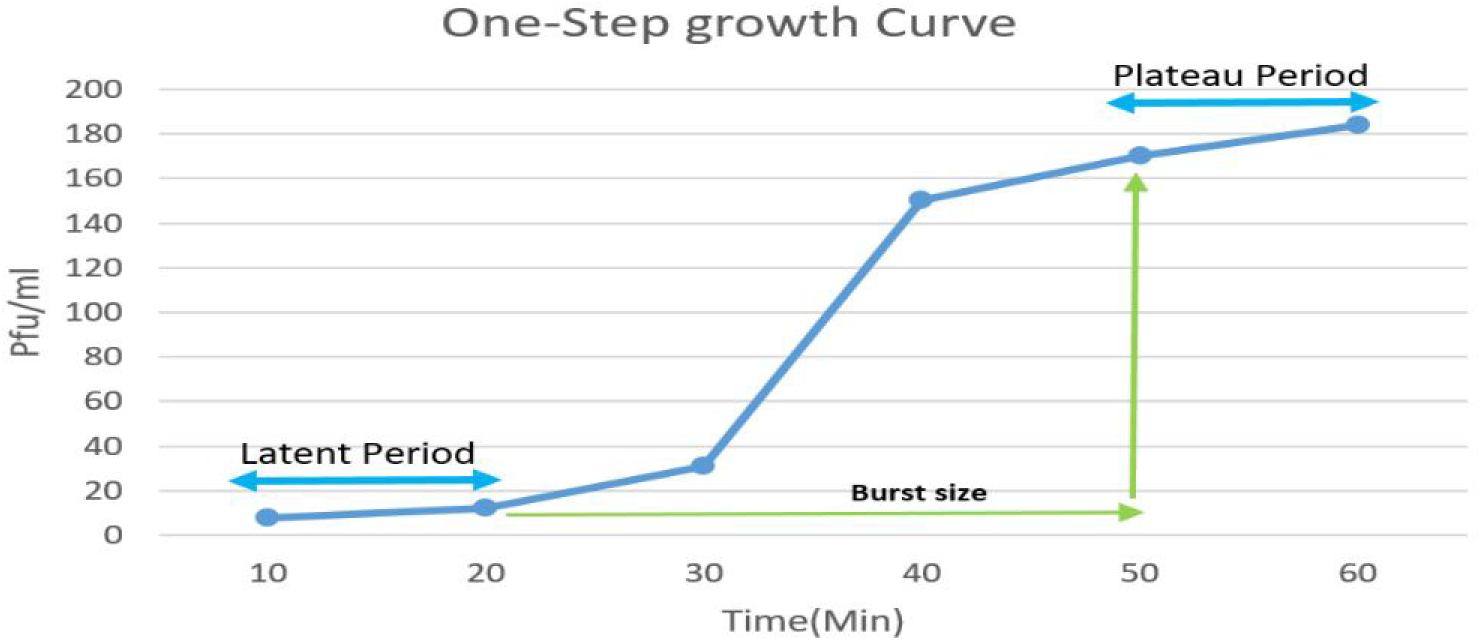
One step growth curve of Phage_KP6697_Omshanti

#### b) Thermal and pH stability

When incubated at various temperatures and pH, the Phage_KP6697_Omshanti survived steadily over a temperature range of 10–80°C.However, the stability of phage significantly decreased when incubated at 70°C for 10 min, decreased to 112 PFU/mL and upon incubation at 80°C for 20 min, the phage completely lost its stability and viability. These results indicated that the phage could tolerate normal temperature (Fig 3). Phage maintained a good activity in the pH range of 5.0–9.0 but its titers dramatically decreased to 1log and 3log at pH 4.0 and 12.0, respectively. When incubated at pH 3.0 and 12.0, the phage was completely inactivated. Thus, the optimum pH range for phage was found to be 5.0–9.0 (Fig 4).

**Fig 3:**
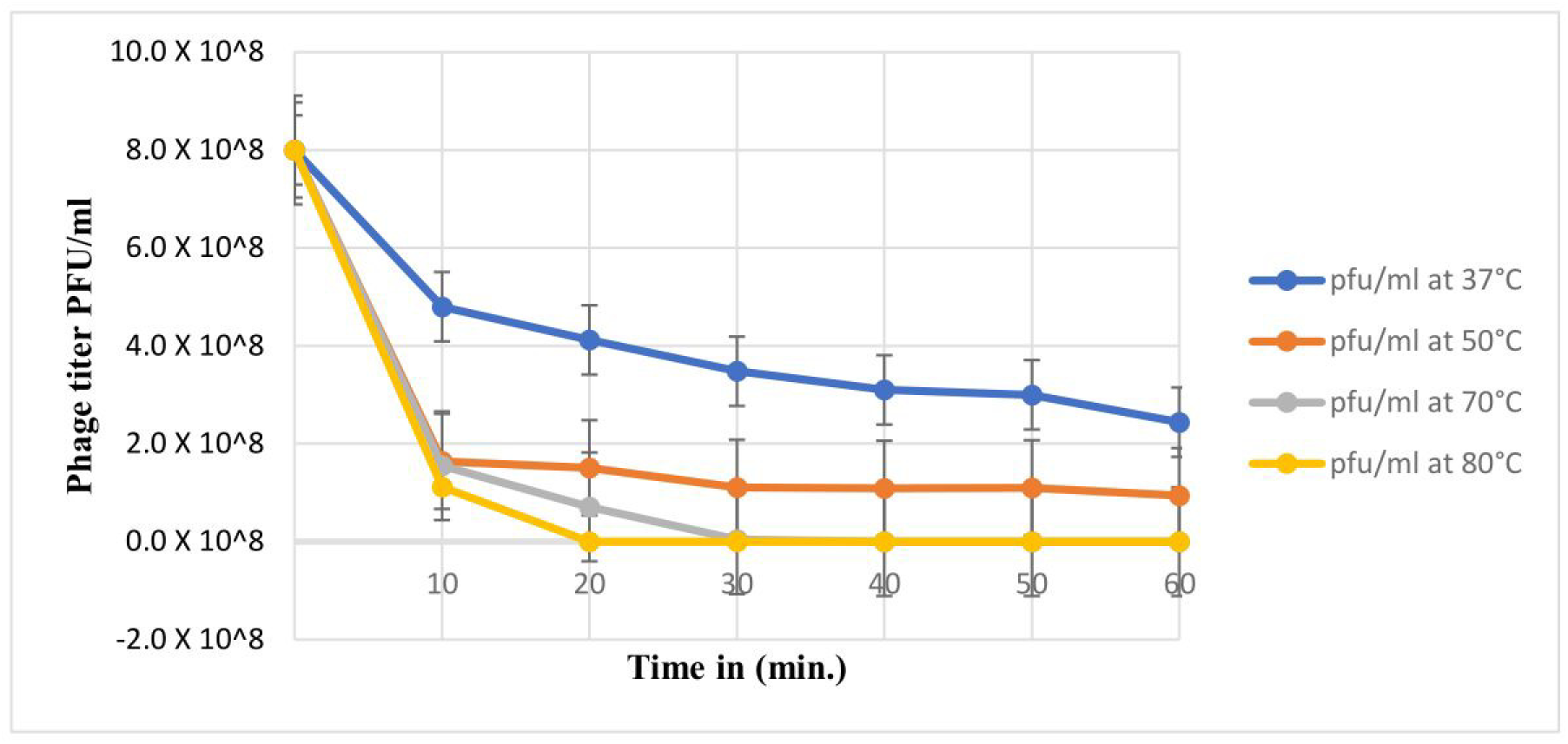
Thermal stability of Phage_KP6697_Omshanti

**Fig 4:**
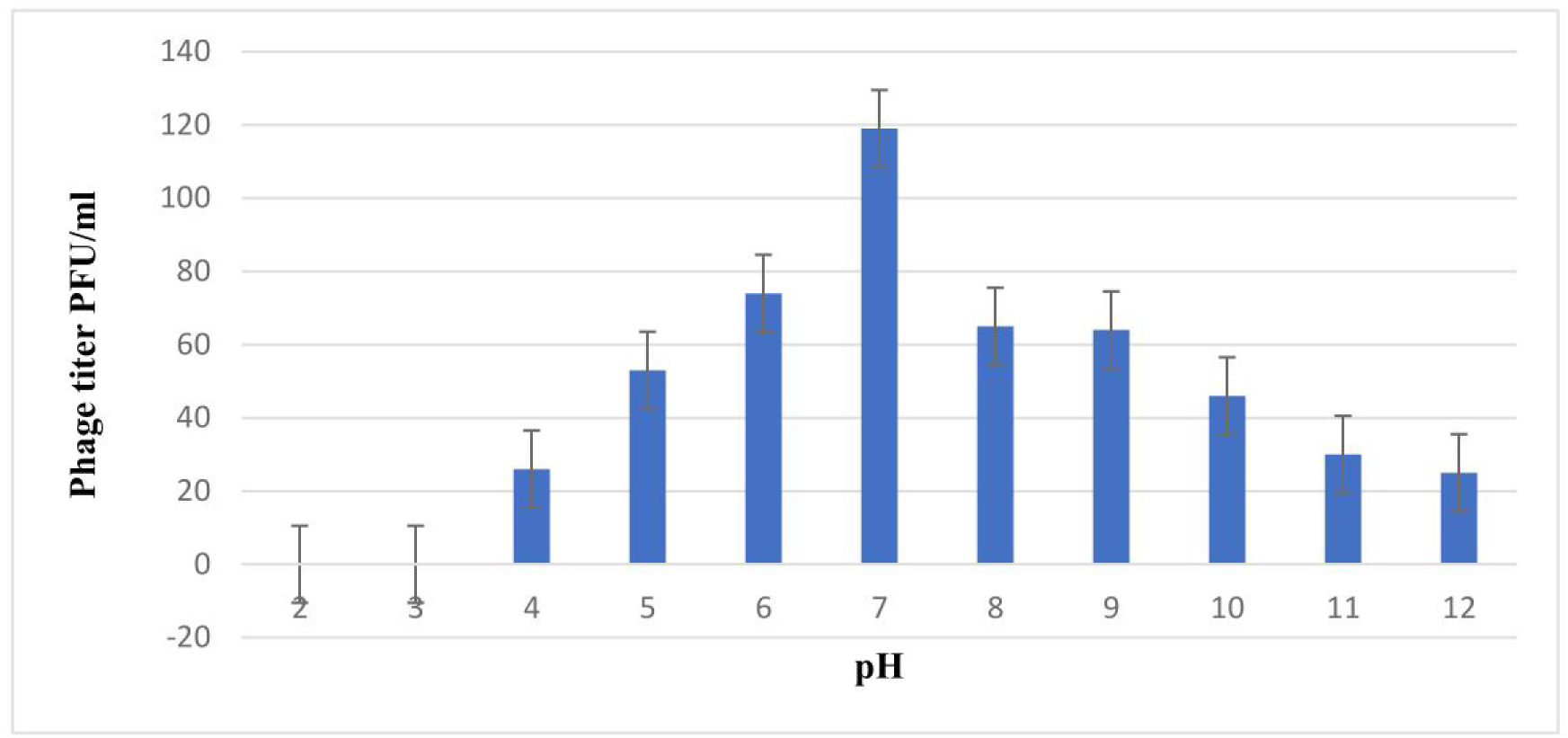
Stability of Phage_KP6697_Omshanti at different pH

#### c) Host range

The Phage_KP6697_Omshanti showed a wide host spectrum on spot assay. A total of 35 strains were tested and 8 were lysed when treated with single phage and 9 were lysed when cocktail of phage was used (Table 3). Single phage mainly lysed 6 species of *Klebsiella pneumoniae*, one of *E. coli* and *Salmonella* Typhi each. However, cocktail of phage lysed 5 strains of *Klebsiella pneumoniae*, 3 *E. coli* and one *Salmonella* Typhi.

**Table 3.**
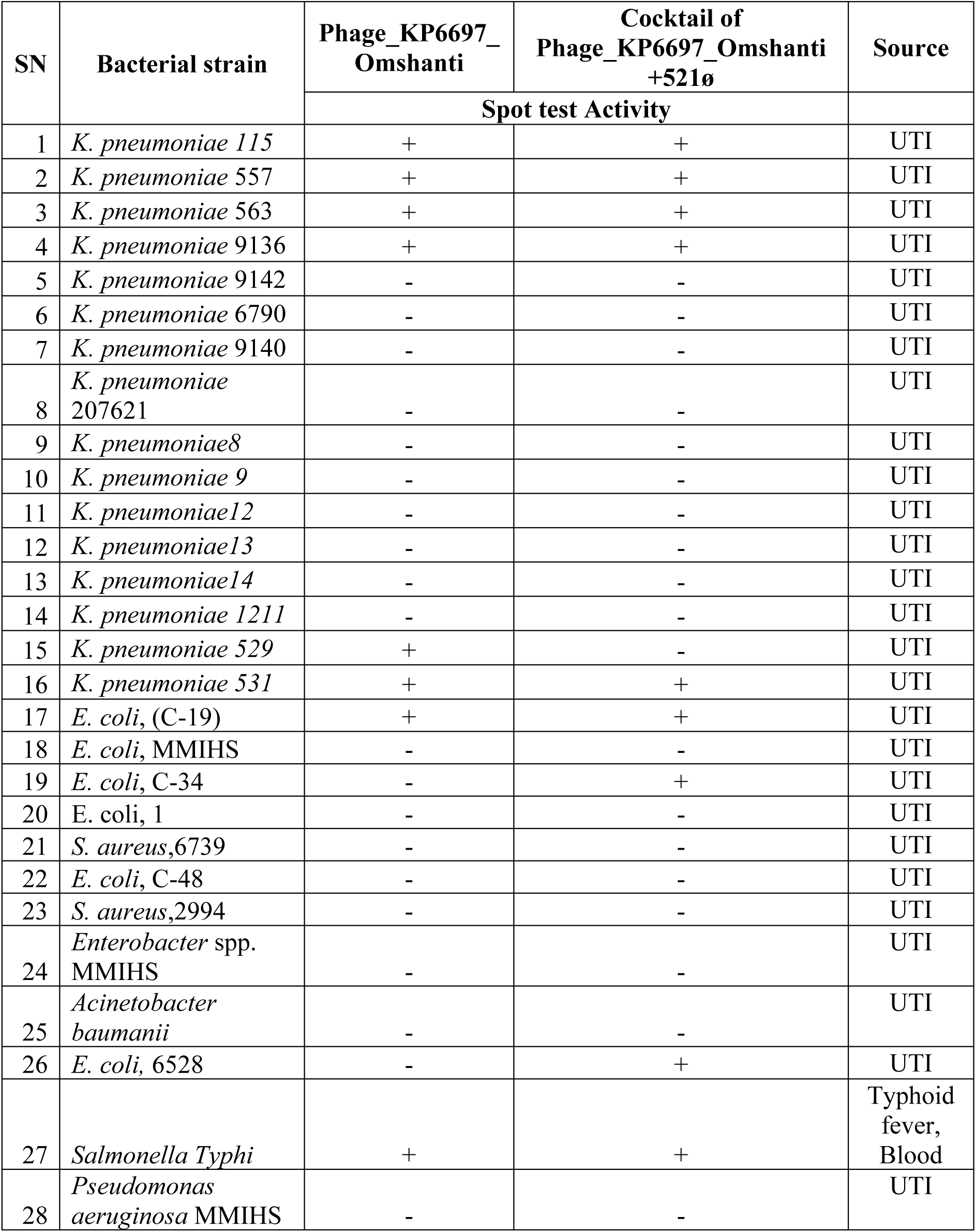
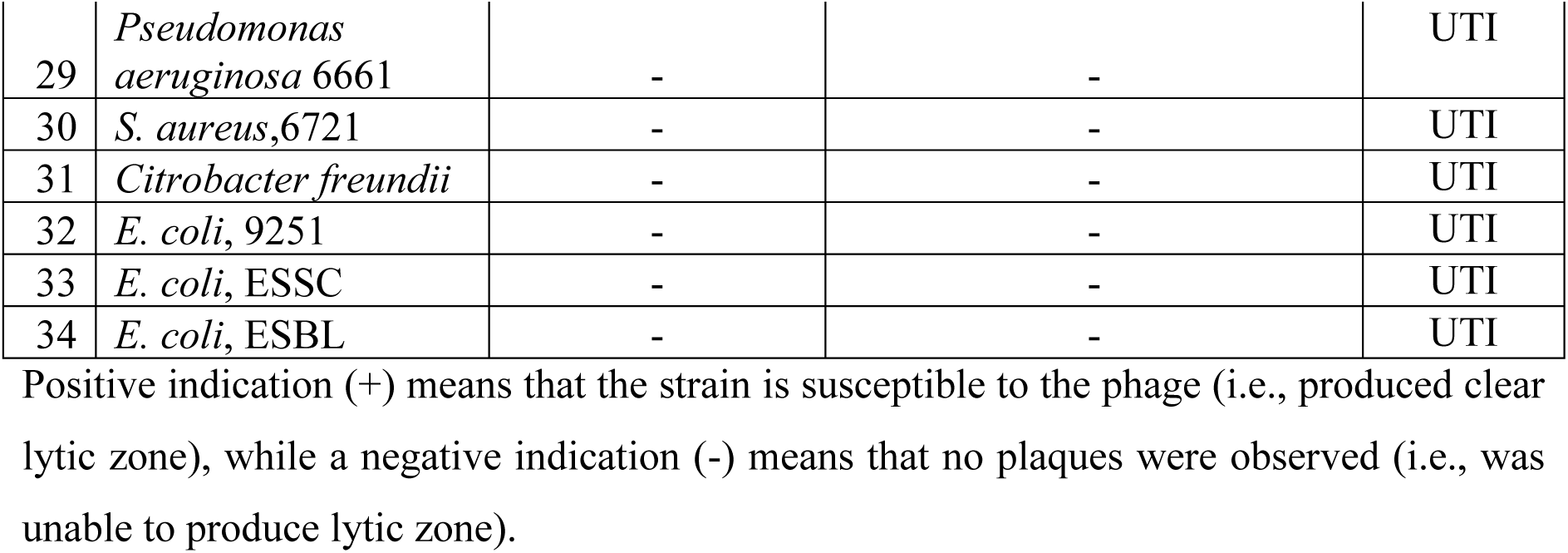
Host range determination for Phage_KP6697_Omshanti.

### Molecular characteristics of Phage_KP6697_Omshanti

Whole genome sequencing revealed that the phage isolated belongs to Caudoviricetes with 45288 bp long genome. The phage genome was compact with few non-coding sections in between and based on the genomic data the phage has virulent(lytic) lifecycle. The complete genome had sequencing depth of 916X with 53.81 % GC content and other details has shown in (Table 4). The phage had total of 54 coding genes out of which 5 genes were associated with lysis, 1 was structural component responsible for packaging, 7 genes were associated with replication, 7 genes were associated with phage assembly, 1 was associated with infection and 13 hypothetical genes were present (Fig 5) (29).

**Fig 5:**
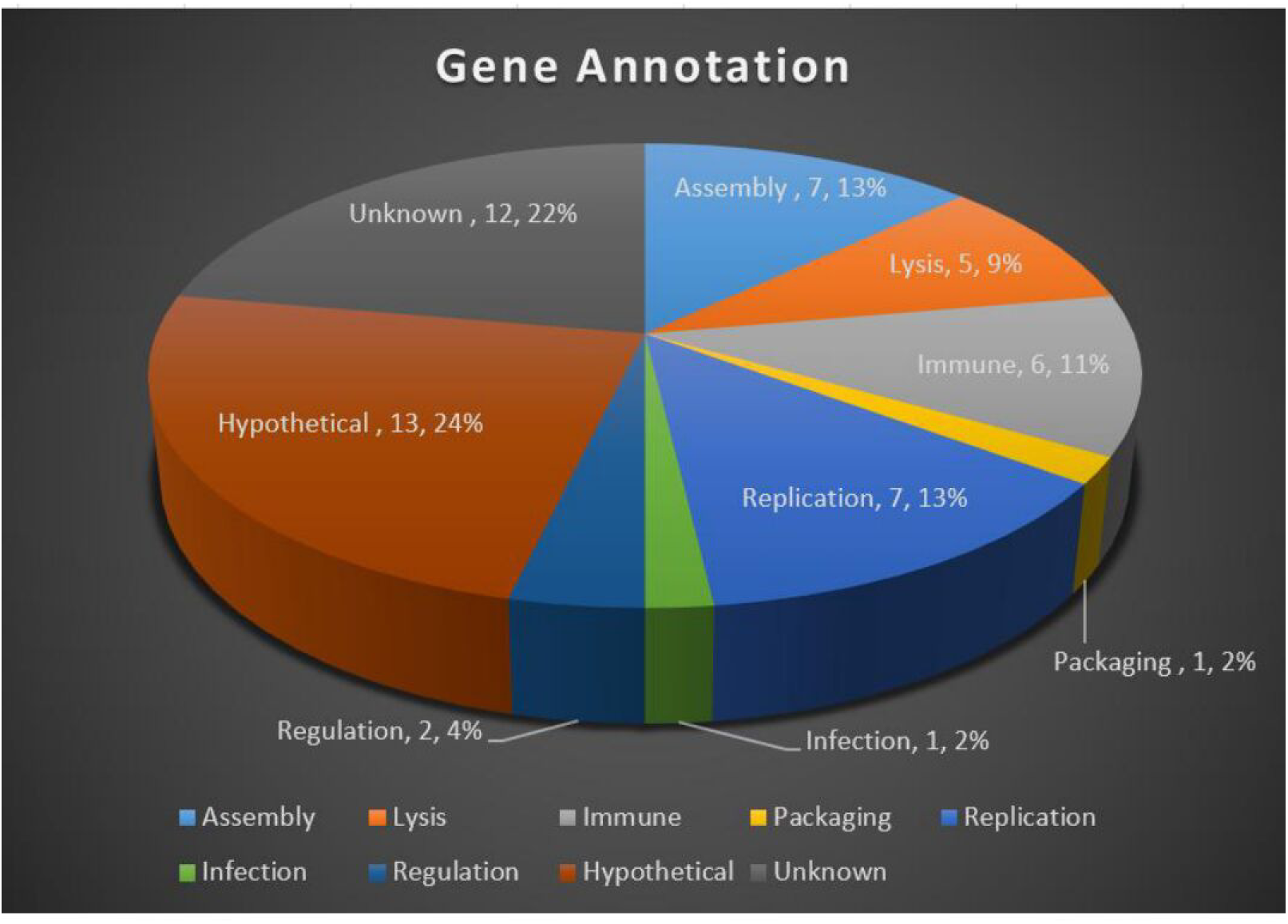
Gene annotation of Phage_KP6697_Omshanti

**Table 4.** Molecular characteristics of Phage_KP6697_Omshanti.

| Phage features | Phage_KP6697_Omshanti |  |  |  |  |
| --- | --- | --- | --- | --- | --- |
|  | Prophage |  |  |  |  |
|  | 153 | 267 | 349 | 529 | 1172 |
| NCBI accession | k141_153 | k141_267_1 | k141_349 | k141_529 | k141_1172_1 |
| **Genomic features** |  |  |  |  |  |
| Length (in base pairs) | 425528 | 40614 | 53577 | 35094 | 39201 |
| Guanine-cytosine (G+C) content | 58.42 | 51.07 | 53.09 | 51.77 | 50.16 |
| tRNAs | 2 |  |  |  |  |
| Average gene size (in bp) | 414 | 55 | 56 | 53 | 41 |
| Completeness | High-quality | High-quality | High-quality | Medium-quality | High-quality |
| Lifestyle | Temperate | Temperate | Temperate | Temperate | Temperate |
| Host | *Klebsiella pneumoniae* | *Klebsiella pneumoniae* | *Klebsiella pneumoniae* | *Escherichia coli* | *Klebsiella pneumoniae* |
| Id | 1 | 1 | 1 | 1 | 1 |
| Taxonomy | – | Caudoviricetes | Caudoviricetes | Caudoviricetes | Caudoviricetes |
| **Protein feature** |  |  |  |  |  |
| Hypothetical proteins | 4 | 10 | 10 | 9 | 6 |
| Assembly gene | 18 | 6 | 2 | 2 | 4 |
| Infection gene | 22 | 3 | 8 | 5 | 8 |
| Assembly and infection gene | 5 | 2 | 5 | 9 | 3 |
| Lysis gene | 20 | 4 | 1 | 0 | 4 |
| Lysis, assembly and infection gene | 1 | 0 | 0 | 0 | 0 |
| Lysis and infection gene | 1 | 0 | 0 | 0 | 0 |
| Lysis and regulation gene | 1 | 0 | 0 | 0 | 0 |
| Lysis and replication gene | 1 | 0 | 0 | 0 | 0 |
| Lysis and assembly gene | 3 | 0 | 0 | 0 | 0 |
| Immune gene | 13 | 0 | 0 | 0 | 2 |
| Replication and packaging gene | 1 | 1 | 0 | 0 | 0 |
| Replication and regulation gene | 3 | 3 | 1 | 0 | 0 |
| Regulation gene | 32 | 2 | 3 | 2 | 2 |
| Regulation and assembly gene | 2 | 0 | 0 | 0 | 0 |
| Regulation, assembly and immune gene | 3 | 0 | 0 | 0 | 0 |
| Regulation and immune gene | 1 | 0 | 0 | 0 | 0 |
| Replication gene | 37 | 1 | 5 | 1 | 1 |
| Replication and immune gene | 1 | 0 | 0 | 0 | 0 |
| Replication and infection gene | 3 | 0 | 0 | 0 | 0 |
| Replication and assembly gene | 2 | 0 | 0 | 0 | 0 |
| Replication, regulation and assembly gene | 1 | 0 | 0 | 0 | 0 |
| Replication and tRNA gene | 1 | 0 | 0 | 0 | 0 |
| Replication and tRNA regulation gene | 1 | 0 | 0 | 0 | 0 |
| Replication, regulation and immune gene | 1 | 0 | 0 | 0 | 0 |
| tRNA and infection | 1 | 0 | 0 | 0 | 0 |
| tRNA | 2 | 0 | 0 | 0 | 0 |
| Integration and assembly gene | 2 | 0 | 0 | 0 | 0 |
| integration, replication, regulation and packaging gene | 3 | 0 | 0 | 0 | 0 |
| Integration gene | 11 | 1 | 0 | 2 | 2 |
| Integration and replication gene | 0 | 1 | 0 | 0 | 0 |
| Packaging gene | 5 | 2 | 6 | 2 | 3 |
| Packaging and assembly gene | 0 | 1 | 0 | 0 | 0 |
| Virulence factors (VFDB database) | 22 | 0 | 0 | 0 | 0 |
| AMR genes (CARD database) | 5 | 0 | 0 | 0 | 0 |

### Functional characteristics of Phage_KP6697_Omshanti

Functional annotation of the phage genome revealed and verified the phage to be virulent phage with lytic lifecycle. The phage sequenced was of medium-size based on genome size (25-100Kbp) (30) containing genes of five different functional categories namely; lysis, replication, packaging, assembly and infection (Fig 6). However, there were some unclassified hypothetical/unsorted genes also present. The genome contained 2 terminator sequences detected in nucleotide location 20569 to 20584 and one in location 28023 to 2804, both the terminator being in the + strand. Furthermore, the genes responsible for replication included proteins such as DNA polymerase family A, DNA helicase along with two DNA-directed RNA polymerases. These proteins are necessary for phage genome replication and transcription of viral proteins. Another functional category i.e., packaging of viral particle included proteins/enzymes such as three different HNH endonuclease, one endonuclease VII, three endonucleases and two different proteins with exonuclease activity along with terminase small subunit protein were present.

**Fig 6:**
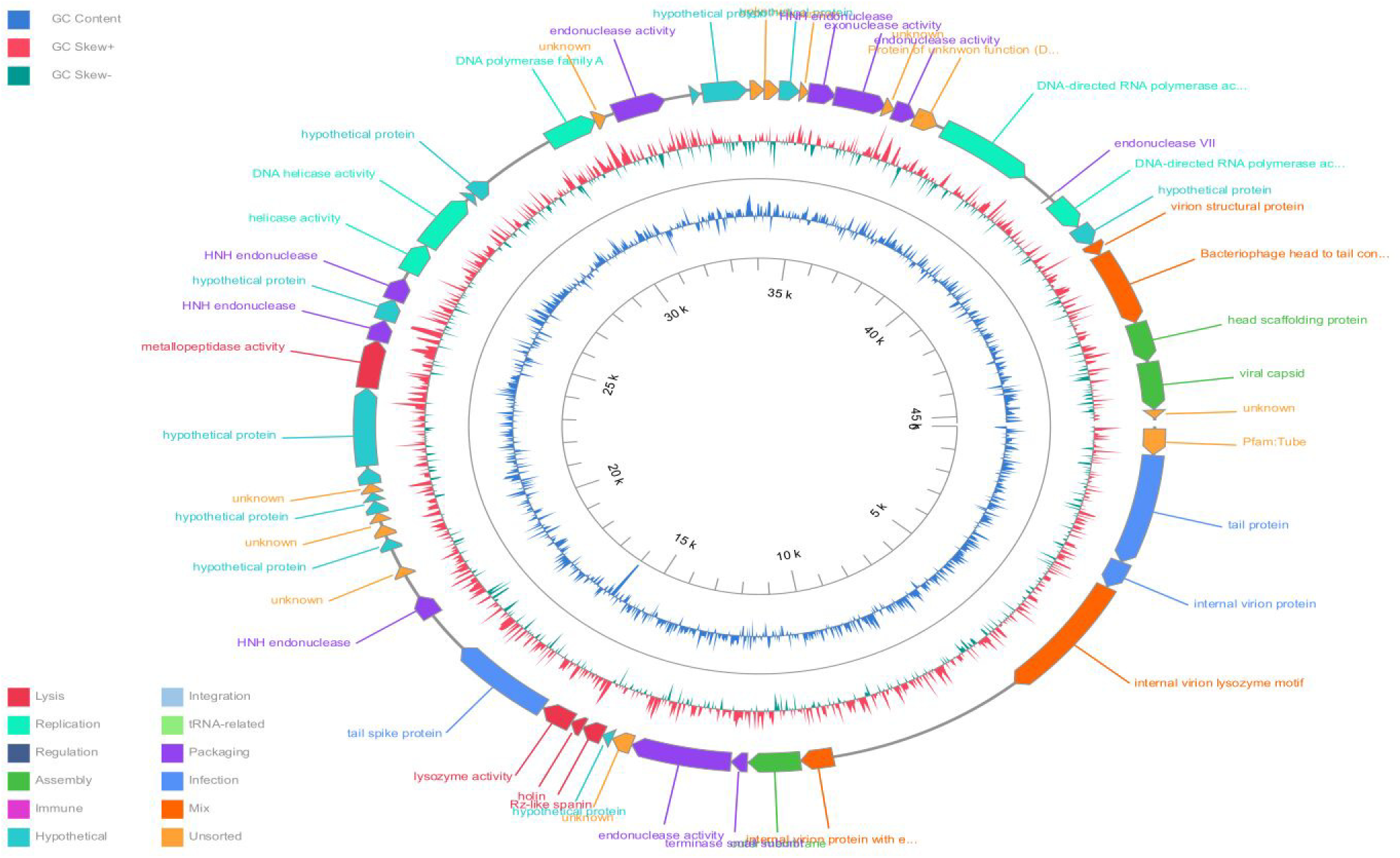
The circular diagram representation of the Phage_KP6697_Omshanti with functional annotation and GC content representation. The image was created from https://phagescope.deepomics.org/

Importantly, structural components that can be used to both infer and validate phage phenotype included head scaffolding protein, viral capsid, virion structural protein, phage head and tail connection and outer membrane proteins. The proteins that are both structural and required for the infection are protein related to tail or tail assembly. Those proteins included; tail protein, internal virion protein (part of tail assembly), tail spike and internal virion lysozyme motif proteins were also present. The most significant functional part of the phage that is needed to lyse the cell for phages to be dispersed is the one classified as functional category of lysis. In this category, classic holin, spanin and lysozyme were present in this phage along with single metallopeptidase like protein. Besides these, the proteins of unknown function and hypothetical protein were also present.

### Effect of Phage_KP6697_Omshanti on biofilm of host KP6697

a. **Optical microscope biofilm imaging**: After 24 h and 48 h of incubation the formation of chains and clusters was observed in control coverslip. While only single cells and small colonies were observed on the experimental coverslip with the addition of single phage (Figs 7 & 8).
b. **Scanning electron microscope (SEM) biofilm imaging:** After 24 h of incubation (biofilm formation by KP6697 bacteria), small microcolonies and filaments were observed on the coverslips incubated without phage treatment (Figs 7 A and C) indicating robust bacterial growth. After incubation for additional 24 h (48 h) the formation of mature biofilm with clusters and channels was observed in culture without phage on the surface of coverslips indicating biofilm formation (Figs 7 B and D). In samples treated with Phage_KP6697_Omshanti alone after incubation for 24 h and subsequent incubation for 48 h, only individual bacterial cells and small aggregates were observed (Figs 8 E-H) indicating inhibition and destruction of bacterial biofilm.

## Discussion

Carbapenem-resistant *Klebsiella pneumoniae* strains have emerged as significant multidrug-resistant bacterial pathogens that pose a serious threat to global public health. Bacteriophages are being considered as a potential therapeutic approach against highly drug-resistant pathogens, including CRKP strains (31). Due to excessive use of antibiotics in hospitalized patients, *K. pneumoniae* exhibits resistance against many antimicrobial drugs. Multiple drug resistant (MDR) *K. pneumoniae* exhibits several mechanisms such as the production of extended spectrum β-lactamases and carbapenemases (32). The MDR strains of *K. pneumoniae* cause different infections in humans, which are difficult to control by using antibiotics. So, bacteriophage therapy is an effective alternative treatment option against multiple drug-resistant pathogens. In this study, a lytic Phage_KP6697_Omshanti against KP6697 bacterium was isolated from a river sample and then characterized for different physiological parameters and the whole genome.

**Fig 7:**
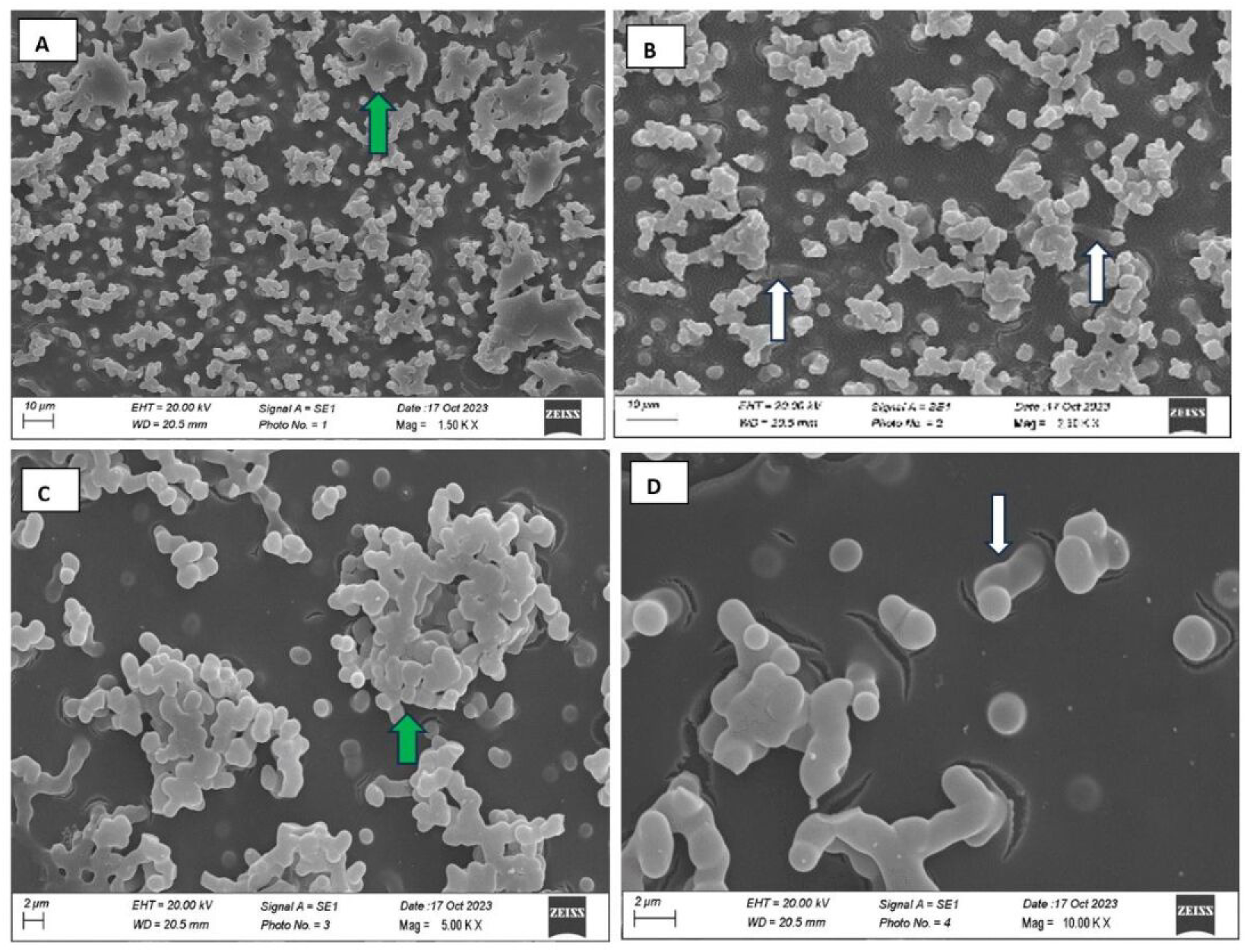
Anti-biofilm activity of Phage_KP6697_Omshanti under SEM (A, B, C, D): After 24 hrs. incubation of *K. pneumoniae* (KP6697) without phages, green arrow marks the formed microcolony, after 48 h of incubation without phages, white arrows indicate the exopolysaccharide connections between bacterial cells, Scanning electron micrographs (SEM). Above, control (not treated with phage), large number of *K. pneumoniae* with intact appearance and high extracellular polysaccharide production.

**Fig 8:**
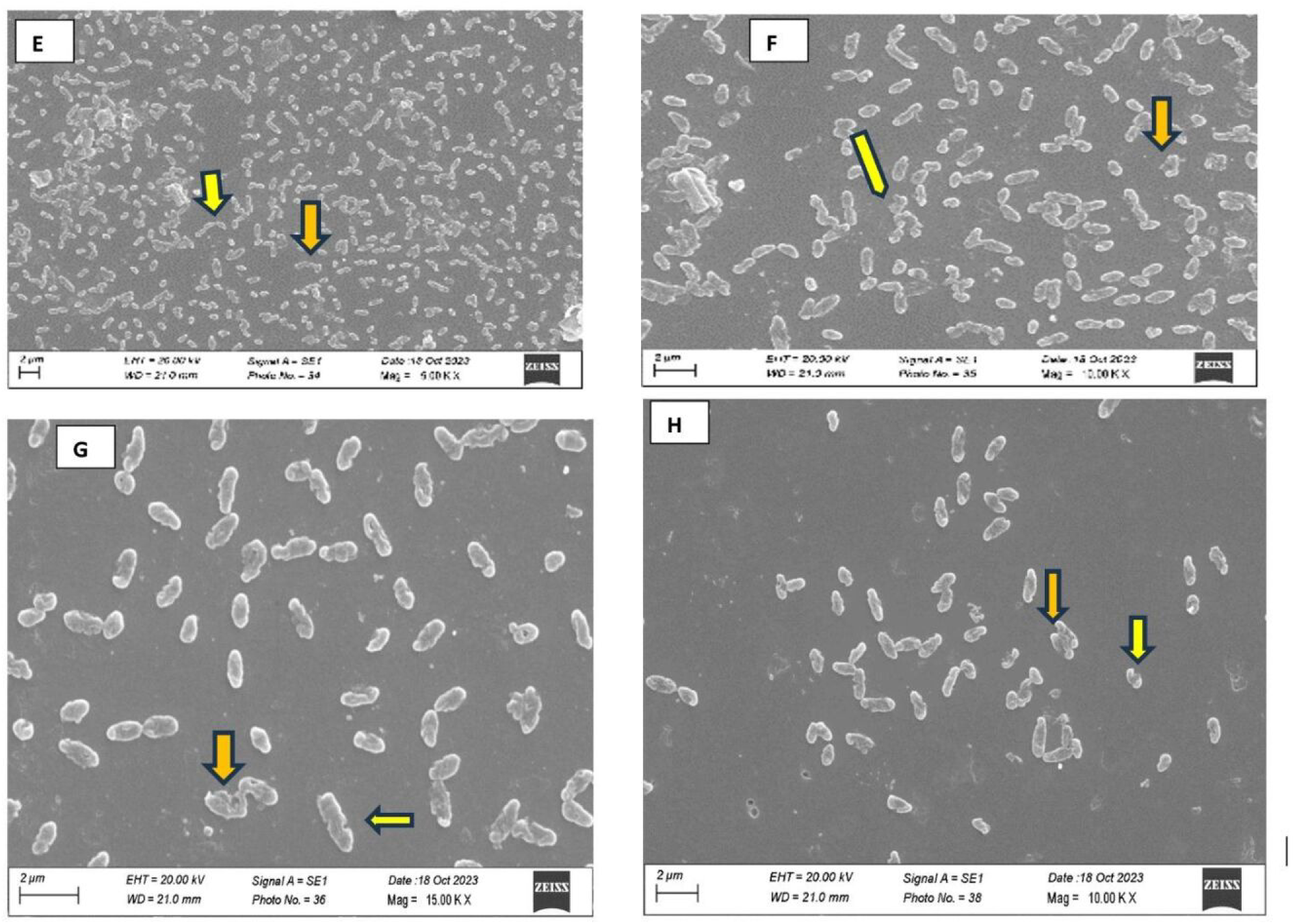
Anti-biofilm activity of Phage_KP6697_Omshanti under SEM (E, F, G, H): After 48hrs. incubation of *K. pneumoniae* (KP6697) treated with Phage_KP6697_Omshanti, showing fewer bacteria presenting cell wall disruption (orange arrow marks), forming depressions (yellow arrow marks) in the configuration of the bacterial skeleton and lessened extracellular matrix

The functional analysis revealed a sophisticated molecular network underlying biofilm formation, with 87 KO entries spanning structural, regulatory, and metabolic categories (Table S1 & S2). Core matrix synthesis machinery was prominent. The complete PNAG pathway (PgaA, PgaB/C, PgaD) indicates active production of this key biofilm component, which enhances structural integrity and antimicrobial tolerance (33). The abundance of PNAG N-deacetylases suggests polysaccharide remodeling to optimize matrix properties (34). Further, Colanic acid biosynthesis (K13650) further supports extracellular matrix formation. Adhesion systems were enriched, particularly type 1 fimbrial genes (K07345 detected six times), underscoring their importance in surface attachment and transition to the sessile state (35). Specialized adhesins such as Mat/Ecp fimbriae (K21967) suggest adaptation to niche-specific colonization (36). Regulatory genes highlighted complex control of biofilm development. Quorum sensing regulators (SdiA, K07782; CsgAB operon regulator, K04333) point to interspecies communication and curli regulation (37),(38) while multiple two-component systems (OmpR, PhoB, CpxR, NarL, GlnG, HydG, GlrR) enable sensing of nutrient limitation, envelope stress, and redox changes (39–42). Nutrient acquisition systems were also central. Redundant siderophore biosynthesis, transport, and receptor genes indicate strong selective pressure for iron, both for growth and competitive advantage (43–45) Secretion systems further support community interactions: a complete Type II pathway facilitates export of enzymes and adhesins (46), and Type VI components (VasG) suggest roles in interbacterial competition(47). Stress response and resistance determinants, including β-lactamase genes (K18699), combined with biofilm-specific barriers such as reduced penetration and persister cells, emphasize the clinical challenge of biofilm infections (48). The integration of adhesion, regulation, and stress pathways highlights the adaptive capacity of biofilms to host defenses and therapy (49–51). Functionally, genes were distributed across matrix synthesis (8.4%), adhesion (12.6%), regulation (16.8%), iron homeostasis (14.7%), secretion (18.9%), cell envelope modification (11.6%), and stress response (17.0%), illustrating a balanced and multifaceted system. Overall, these findings underscore the complexity of biofilm development, revealing potential intervention targets such as PNAG synthesis, quorum sensing, siderophore uptake, and secretion systems. The identified pathways highlight why biofilms are central to antimicrobial resistance, chronic infections, and environmental persistence, while also guiding strategies for anti-biofilm therapy.

We tested several samples including Balkhu river sample in Kathmandu valley for the presence of lytic phage against KP6697 as a bacterial host. We could find only one lytic bacteriophage against *Klebsiella pneumoniae*. Our results indicated that the host ranges of our phage appeared to be narrow and were specific to certain *K. pneumoniae* strains. Interestingly, we identified a phage with a relatively wide intra specific host range analysis. Phage_KP6697_Omshanti was examined for its lytic ability against 16 strains belonging to *K. pneumoniae* and other unrelated genera. It lysed 6 of 16 (37.5%) CRKP strains from our collection while it showed lytic activity against 18 tested members of other genera, out of them, 2 were lysed (n=18, 11.1%) in MDR interspecies strains, depicting its narrow host range (Table 3) But our isolated phage seemed to be effective against *E. coli* and *Salmonella* Typhi host bacteria other than *Klebsiella pneumoniae*. These results demonstrate that Phage_KP6697_Omshanti showed host specificity within the genus *Klebsiella* but lytic activity against two inter genus also. Previously, the bacteriophages KP1513 and KP-34 against *K. pneumoniae* also showed narrow host range (52, 53). No significant result was obtained for inter specific host range analysis. TEM of our phage revealed that phage belonged to the class Caudoviricetes of the order Caudovirales.

Both temperature and pH are crucial factors that influence the bacteriophage stability. Temperature affects the whole phage replication process and also regulate phage viability, occurrence and storage (54). Our phage showed the highest activity at 37°C while a reduction in phage titer was observed at all other temperatures (Fig 3). The phage was completely inactivated at 70°C temperature, almost similar to Kp34virus genus phages as described earlier (52). It showed more activity at alkaline pH than acidic pH (Fig 4), however, 7.0 pH found optimal for this phage. Due to its stability at alkaline pH, it can be used as a therapeutic agent for *K. pneumoniae* mediated urinary tract infection (UTI) and infected wounds. It can be used in the impregnation of urinary catheters to inhibit bacteriological biofilm as previously suggested (55). We studied the growth parameters of Phage_KP6697_Omshanti through a single-step growth curve which helps in defining phage lytic potential for biocontrol of bacteria (56, 57). Our results indicate a short latency period of approximately 20 min. and a considerable burst size of approximately 119 virion per bacterium. While the latency period defines the period between virion attachment to the host bacterium and the release of new phage particles, burst size defines the average number of new phage particles released from each infected cell after a lytic cycle (58). Therefore, short latency and large burst size are considered characteristic features of an efficient lytic phage and indicate the suitability of a phage for therapeutic applications (58, 59).

The Phage_KP6697_omshanti genome encodes a sophisticated array of genes that collectively mediate bacterial lysis, biofilm disruption, evasion of host defenses and efficient progeny production. Key lysis genes such as holins, endolysins, Rz-like spanins and metallopeptidases work synergistically to breach the bacterial cell envelope by forming pores, degrading peptidoglycan and disrupting the outer membrane, facilitating virion release (Table 4) & (Fig 5). Beyond direct lysis, phage tail spike proteins target bacterial surface structures to penetrate and destabilize biofilms, aided by enzymatic degradation of the extracellular matrix. The phage also encodes multiple nucleases that counteract bacterial restriction-modification systems and degrade regulatory DNA involved in biofilm maintenance, while helicase and DNA polymerase activities ensure rapid phage genome replication overcoming host repair mechanisms. Structural and packaging proteins further support effective virion assembly and DNA encapsidation, enabling sustained infection cycles in biofilm-protected bacteria. Together, these genetic components highlight the phage’s evolved capacity to lyse bacterial hosts and disrupt biofilms, underscoring its potential as a therapeutic tool against biofilm-associated infections (60–62).

In many studies of the effects of bacteriophages on biofilms, no difference in efficacy is observed between the use of a single phage and a cocktail of bacteriophages. For example, a study of the phages 39APmC32, 65APm2833 and 72APm5211 showed that the antibiofilm activity of phage cocktails on 24hrs. *Proteus mirabilis* biofilms was similar than that of the most active phage (63). In another study, the pretreatment of 48hrs. old biofilm with the *Pseudomonas aeruginosa* phage cocktail resulted in a titer reduction of the same order of magnitude as with a single phage application. However, researchers reported that the application of the phage cocktail rather than single phages prevented the appearance of phage-resistant bacteria (64).

Thus, in the present study we decided to perform the efficacy of single phage in biofilm disruption. The present study demonstrated the ability of single phage with depolymerase activity to effectively control antibiotic resistant *K. pneumoniae* biofilms. In control of the experiment, all stages of biofilm formation were observed, from single adherent cells, formation of small colonies, clusters and to biofilm formation and maturation. It is noteworthy that filament formation by *K. pneumoniae* cells was observed in the present study. The production of filaments has been associated with the process of biofilm formation and modern research indicates that filamentation plays a vital role in biofilm development (65). There is also evidence that exposure to beta-lactam antibiotics is associated with an increased frequency of filament formation (66), it has been reported that *K. pneumoniae* cells could form filaments in response to cefotiam and cefazolin (67). We hypothesize that since the biofilm is composed of heterogeneous cells with different metabolism which plays a key role in adaptation (68, 69). It is possible that exposure to antibiotics leads to the selection of filamentous cells that are already presented in the population.

Bacteriophages were added after 24hrs. of biofilm development, when such stages of biofilm growth as micro-colony and filament formation were observed in the control samples. These time points were chosen based on previous studies (70, 71) and to monitor the effect of bacteriophages on the transition to more mature stages of bio-film formation. The addition of bacteriophages to the cell culture prevented biofilm formation; only single cells and small microcolonies were observed in the samples with phages.

The SEM result suggested that Phage_KP6697_Omshanti can not only inhibit biofilm formation but also degrade the formed mature biofilms. Many bacteriophages produced specific depolymerases which are able to destroy the capsular polysaccharides, thereby allowing adsorption of phages to the outer membrane receptors (72) and penetration of phage DNA into the bacterial cells (73).These phage-derived capsule depolymerases can be potentially used as therapeutic agents against capsulated bacteria such as CRKP strains. In fact, besides depolymerases, phage-encoded lysozymes (endolysins, lysins, etc.) can also digest bacterial cell wall material (74, 75). Lysin gene were found in the genome of the phage based on BLASTn predication. Therefore, we suspected that the mechanism of biofilm formation inhibition and biofilm degradation may be associated with holin and endolysin gene of Phage_KP6697_Omshanti. The results suggested that Phage_KP6697_Omshanti could be an attractive candidate as an anti-CRKP agent. The biofilm removal tendency was not different for biofilms of different ages as the difference in biomass reduction was not statistically significant (76). In this study, complete eradication of the bacterial culture was not achieved, but a significant decrease in number of bacteria and biofilm destruction was observed (Figs 7 & 8). However, combined phage-antibiotic therapy may be a solution for complete eradication of bacterial colonization when necessary. A combination of a pre-adapted bacteriophage and antibiotic is known to be used in the treatment of a patient with fracture-related pan drug resistant *K. pneumoniae* infection (77). However, Complete lysis was not observed, but we presume that this is extremely difficult to achieve with bacteriophage treatment, since even between lytic phages and bacteria more complex mechanisms that regulate lysis exist, such as quorum sensing, hibernation or transient resistance (78–80).Researchers also observed inherently susceptible but phenotypically antibiotic-resistant subpopulations of bacteria (81), the presence of persisters in the bacterial population can potentially influence phage infection as well. Moreover, complete elimination may often not be necessary even in the medical treatment since in humans *Klebsiella* spp. can be found as commensals of the gastrointestinal tract, mouth and nasopharynx (82), thus a significant reduction in number, biofilm disruption and restoration of normal microbiota balance can become clinically significant. The combined phage-antibiotic therapy could be a solution for the complete eradication of biofilms and bacterial colonization in those cases where it is necessary (77, 83).

The viral RNA polymerases are central to phage biology and biotechnology applications particularly in mRNA-based therapies due to their efficient transcripition capabilities. Notably, certain phages like T7 encode their own single subunit RNA polymerases (RNAPs) which selectively transcribe viral genes without relying on host machinery (84, 85).Some phages modify host RNAPs through covalent modifications or accessory proteins to redirect activity towards viral transcription (86). Bacteriophage derived RNAPs from T7 and SP6 have been instrumental in large scale mRNA-synthesis for vaccines and therapeutics (85) due to their high processivity and specificity. These enzymes show evolutionary relationships with other nucleotide polymerases suggesting broader functional diversity. Recent findings highlight that some jumbo bacteriophages encode multi-subunit RNA polymerases similar to those found in cellular organisms (87) offering new strategies for controlling gene expression during infection. Collectively, these enzymes enable rapid virion production by ensuring efficient genome duplication and controlled gene expression within host cells. Understanding these mechanisms provides insights into phage host interactions at a molecular level which can be leveraged for developing novel therapeutic strategies using phage therapy against bacterial infections.

Understanding the structural components of bacteriophages is crucial for inferring and validating their phenotypic characteristics. In this study, we identified key structural proteins that contribute to both virion architecture and infection mechanism. The head scaffolding protein, viral capsid, virion structural protein, phage head tail connection and outer membrane proteins are essential components that dictate the structural integrity and stability of the phage. These components play a significant role in determining the morphology and assembly of the virion thereby influencing phage classification and host specificity (88, 89). Among the structural proteins, those related to the tail and tail assembly are particularly important for infection. These proteins include tail proteins, internal virion proteins involved in tail assembly, tail spikes and internal virion lysozyme motif proteins. The tail proteins mediate initial host recognition and attachment, a crucial step in the phage life cycle (90). Tail spikes often facilitate enzymatic degradation of bacterial surface components, enhancing the efficiency of host invasion (91). Internal virion proteins, as part of the tail assembly, contribute to the structural organization required for successful DNA injection into the host cell (92). Furthermore, lysozyme motif proteins play a pivotal role in breaching the bacterial cell wall enabling genome delivery and initiating the infection process (93). The presence of these structural and infection related proteins highlights their dual role in both maintaining virion architecture and ensuring effective host interaction. By analyzing the conservation and variation of these proteins among different phage families, we can refine our understanding of phage evolutionary relationships and host adaptation strategies (94). Additionally, these findings provide a framework for engineering phages with enhanced therapeutic potential particularly in the development of phage therapy against antibiotic resistant bacteria. Future studies should focus on the functional characterization of these proteins to elucidate their specific roles in infection dynamics and their potential applications in biotechnology and medicine.

## LIMITATIONS

This study did not determine the exact mechanism by which Phage_KP6697_Omshanti lyses its host bacteria, as it was beyond the project’s scope. Further research is needed to understand its inter-genus lysis capability which could aid in designing synthetic phages with broader host ranges. Limitations include testing the phage only against CRKP isolates from a single hospital and the full characterization of only one phage isolate.

## CONCLUSIONS

This study highlights the urgent threat posed by carbapenem-resistant *K. pneumoniae* (CRKP), which combines multidrug resistance with robust biofilm formation, complicating treatment and persistence in clinical and environmental settings. The functional analysis of KO genes revealed a complex biofilm-associated network encompassing adhesion, matrix synthesis, regulatory circuits, secretion, and stress response pathways, underscoring the adaptive resilience of CRKP. Against this background, we characterized Phage_KP6697_Omshanti, a lytic bacteriophage with strong activity against CRKP and some interspecies MDR strains. The phage demonstrated stability under physiologically relevant conditions, a short latency period, and high burst size, all features favorable for therapeutic use. Genomic analysis further revealed lysis and biofilm-targeting genes, including depolymerases and endolysins, supporting its potential as an anti-biofilm agent. Importantly, SEM observations confirmed its ability to inhibit and degrade biofilms at different maturation stages.

Although complete eradication of bacterial populations was not achieved, significant reductions in biofilm biomass were observed, suggesting that phage therapy, alone or in combination with antibiotics, may provide clinically meaningful outcomes. The structural and enzymatic components of Phage_KP6697_Omshanti not only reinforce its therapeutic potential but also expand opportunities for biotechnological applications, such as engineered phages or phage-derived enzymes for combating MDR pathogens. Collectively, these findings position Phage_KP6697_Omshanti as a promising candidate for developing targeted therapies against CRKP infections. Future work integrating phage–antibiotic synergy, functional characterization of structural and regulatory proteins, and evaluation in in vivo models will be critical for translating these insights into effective therapeutic strategies.

## MATERIALS AND METHODS

### Bacterial strain and culture conditions

All *K. pneumoniae* used either as host (N=1) or host range spectrum analysis (N=35) were received from the Sukra Raj Tropical and Infectious Diseases Hospital, Teku, Kathmandu, Nepal. Preliminary identification of the *Klebsiella pneumoniae* (KP6697) bacteria was performed via polymerase chain reaction (PCR) amplification of the 16S rRNA gene using universal primers and protocols (95). Whole genome sequencing of this bacterium was also performed by using the Illumina (San Diego, CA, USA) MiSeq platform. This bacterium was cultured and maintained in Luria–Bertani (LB) medium at 37°C on an orbital shaker at 180 rpm. Phosphate-buffered saline (0.1M Na_2_HPO_4_, 0.15 M NaCl_2_, pH 7.2) was used for dilution and washing of bacterial cells unless specified otherwise.

### Antimicrobial susceptibility testing

Antimicrobial susceptibility testing (AST) was conducted using fully automated Vitek2 Compact System-(bioMeriux, France) following the Clinical & Laboratory Standards Institute (CLSI) guidelines. Meropenem and imipenem showed ≥16µg/mL. The 2023 CLSI breakpoints were used to interpret the susceptibility results (96). All the AST experiment were performed in three biological replicates.

### Phenotypic detection for MBL -β-lactamase

The initial screening test for the production of MBL (metallo-beta-lactamase) was performed by using imipenem (IMP, 30μg) disk (Mast, UK). If the zone of inhibition (ZOI) was ≤27mm for imipenem, the isolate was considered as a potential MBL producer. The organism was swabbed on to a Mueller-Hinton agar (MHA, Hi-Media, Mumbai, India) plate as done for the screening test in the antibiotic sensitivity test. Then, the combination disk (CD) method was applied for the confirmation of MBL producing strains according to the following protocol (97).

### Combination disk (CD)method

Combination disk methods were used for the confirmation of MBL-producing strains in which IMP alone and in combination with EDTA were used. An increased ZOI of ≥5mm for either antimicrobial agent in combination with EDTA versus its zone when tested alone confirmed MBL production (98). *K. pneumoniae* ATCC BAA-1706 (1706, negative for carbapenemase production), *K. pneumoniae* ATCC BAA-1705 (1705; serine enzyme (KPC) positive) and *K. pneumoniae* ATCC BAA-2146 (2146; MBL (NDM) enzyme positive) were used for quality control (96).

### Bacterial host DNA extraction, sequencing, and genome analysis

The genome of host bacteria KP6697 was extracted utilizing Cetyltrimethylammonium Bromide (CTAB) method according to established protocol as described elsewhere (99). Whole-genome sequencing was performed on Illumina Miseq platform using DNA prep kit (20060060). The genome sequences were assembled using Spades genome assembler v 4.2 with –careful flag. The contigs were then subjected to taxonomic classification with Kraken2 (100) for species identification along with checkVv1.0.3 (101) for identification and quality assessment of a viral contigs and prophage detection. Following species identification, the reads were mapped against reference sequence (CP052388.1 *Klebsiella pneumoniae* strain C17KP0052) with Bwa-mem2 v 2.3 (102) and consensus was generated with samtools mpileup (103) followed by Vcftool-consensus (104). The consensus was scanned against pubMLST using GalaxyAustralia (https://usegalaxy.org.au/). The whole genome was annotated with bakta v1.11.2 and functional annotation was carried out with BlastKOALA (105). Furthermore, social gene responsible for biofilm formation and quorum sensing were identified with SOCfinder v1.4. (106). The plasmid and AMR the contigs were analyzed with staramr that scans genome assemblies against the ResFinder and PlasmidFinder (107, 108) databases searching for AMR genes in GalaxyAustralia.

### Phage isolation and purification

Water samples were collected from a Balkhu river of Kathmandu valley in a sterile 50.0 mL Falcon tube between July and September 2022. Before collection, the water was mixed thoroughly, and the sediments were collected with the overlying water from collection sites. Phage isolation was performed by soft agar overlay technique as described previously (109). A completely isolated clear plaques were picked by using sterile tooth tip and dissolved in 1.0 mL sodium chloride-magnesium sulfate (SM) buffer (50 mM Tris−HCl [pH 7.5], 100 mM NaCl, 10 mM MgSO_4_, and 0.01% gelatin). The mixture was filtered through 0.2μm syringe filter (Axiva Sichem, Haryana, India) to remove the bacterial contamination. The filtrate was further used for soft agar overlay assay and next day, an isolated plaque was picked. The process was repeated 3 times and the pure phage strain was collected from the plates of last round. For this, the plates from third round containing plaques were flooded with 10.0 mL of SM buffer and 2 drops of chloroform was added into it. The plates were sealed and incubated at rotating shaker (80 rpm for 30 minutes) for phage elution/diffusion from the plaques. The SM buffer was then collected in a Falcon tube and centrifuged at 4000 rpm for 20 min. Then, the supernatant was filtered through 0.2μm syringe filter (Axiva Sichem, Haryana, India) to obtain high titer of pure phage strain. The purification, counting, and propagation of phage was performed using the double-layer agar plate method as described elsewhere (110). SM buffer was used for the dilution of the phages. Finally, the phage lysates were stored at either 4^◦^C or at -80^◦^C in glycerol (3:1 [v/v]) (111) until further use. All the sterile operations were carried out in class II biosafety cabinet with institutional safety approval.

### Transmission electron microscopy

The phage lysate was fixed with 2% paraformaldehyde and 2.5% glutaraldehyde. Ten microliter phage lysate was spread on a carbon-coated copper grid and negatively stained with 2% (w/v) uranyl acetate (pH 4.5). The copper grid was dried and examined under the FEI Tecnai T-12 Transmission Electron Microscope (TEM) (FEI Company, Hillsboro, OR, USA) at an accelerating voltage of 80Kv. Bacteriophage morphology and taxonomy were confirmed following the guidelines from the International Committee on Taxonomy of Viruses (112).

### One-step growth curve and burst size determination

A single-step growth curve was performed following Kim *et al*. (113) to determine the phage latent period and burst size. In brief, 10 mL exponentially growing host bacteria KP6697(OD=0.08-0.1 at 600nm) was infected with purified phage particles at an MOI of 0.1 and was allowed to adsorb for 15 min at 37°C. Subsequently, cells were pelleted by centrifugation (12,000×g for 5 min) and unadsorbed phages were removed by washing the pellets with fresh TSB. Cell pellets were then resuspended in 10 mL fresh TSB and incubated at 37°C. Cultures were incubated for 120 min and after every 10 min, a sample 100µL was taken for double layer agar plaque assay. Each experiment was conducted in triplicate.

### Thermal and pH stability

The stability of isolated phage at different temperatures and pH was determined according to D’Andrea *et al* (114) with slight modification. Briefly, known phage lysate (10^8^PFU/mL) in SM buffer was adjusted to different pH ranging from 2 to 12. Phage suspensions were incubated for 60 minutes at 37°C and then titrated using a double-layer agar assay on the primary host. For temperature stability, known phage lysate (10^8^ PFU/mL) was aliquoted into the Eppendorf tube and incubated at (37°C, 50°C, 70°C and 80°C) for up to 180 minutes and titrated by using double-layer agar assay as described earlier.

### Host range analysis

Purified phage was tested for their host range on 34 different multidrug-resistant clinical isolates. Initially, spot assay was done to determine the host range of the phages as described previously (115). Based on the spot assay results, the lysis efficacy of the phages on bacterial strains was assessed by Efficiency of Plaquing (EoP). For EoP, 100μL of an overnight culture of each isolate was mixed individually with 3.0 mL of semi-solid top agar (TSA, agar = 0.4%, temperature = 50°C) and immediately poured onto bottom agar plates (TSA, agar = 1.0%). The top agar was allowed to solidify at room temperature. Then, 10.0μL of 10-fold serial dilutions of the phage lysate (10^8^ PFU/mL) were spotted on the bacterial lawn and dried until completely absorbed on the top agar. The plates were then incubated overnight at 37°C. After incubation, the lowest titer of the phage that gave countable plaques was determined and the phage titer was calculated. The EoP was determined by dividing the average number of plaques formed on the tested bacterial strain by the average number of plaques on the original host bacterium. The test was performed in triplicates.

### Anti-biofilm activity of the phage

Biofilm of host bacteria KP6697 was established in 8-well polystyrene tissue culture microplates to achieve an improved cell attachment. TSB supplemented with 1% D-(+)-glucose was used to perform this assay, as this helps to improve biofilm formation (116). Briefly, an overnight culture was in the microplate wells a 1:20 dilution was performed by adding 10µL of the bacterial suspension to 190µL of TSB. Two hundred microliters of broth were added to a set of wells as a negative control. All wells were replicated three times. Afterwards, microplates were incubated at 37°C for 48hrs. with no shaking for biofilm formation. During the incubation time (24hrs. after incubation). 100µL of fresh glucose was added to all control and test wells. Following incubation, medium was poured off and wells were carefully washed twice with sterile phosphate-buffered saline (PBS) solution to remove any planktonic cells. Micro-plates were allowed to dry for 1hr. at 50°C. To determine total biofilm biomass, microplate wells were stained with 0.1% crystal violet (CV). After staining, the wells were washed twice with PBS solution and dried. Biofilm formation was determined by visual comparison of the stained wells and photographed. For optical density readings of the staining intensity, 10µL of 95% (v/v) ethanol was added to each well and optical density at 590 nm (OD590) was measured using a plate reader. After 24hrs. of biofilm formation, 0.1 mL of phage with a concentration of 10^8^ PFU/mL of a single phage was added in 8-well polystyrene tissue culture microplates and incubated for 48hrs. at 37°C. Glass coverslips were added in all wells with and without phage addition. For optical microscope visualization, coverslips were removed with tweezers and placed into petri dishes with paper filters at the bottom without drying. To preserve the natural form of biofilms, samples were vapor-fixed with 25% glutaric aldehyde for 3hr. After fixation, the samples were stained with DAPI (Sigma-Aldrich, Germany). The effect of phage on biofilm formation and destruction was evaluated using an AxioImager A1 light microscope (Carl Zeiss, Germany). After 24hr. of biofilm formation, a single phage was added to the biofilm and incubated for additional 24hrs. Control were also incubated for 72hrs. without the addition of phage.

### Scanning electron microscopy (SEM)

Few samples of biofilm producing host bacteria KP6697 treated with phage were fixed with 2.5% glutaraldehyde for 60 min, then dehydrated in a graded ethanol series (30, 50, 70, 80, and 96%) and placed into acetone. The samples were dried in a critical-point dryer HCP-2 (Hitachi Ltd., Japan) and coated with Au–Pd in IB-3 ion coater (Eiko Engineering Co., Tokyo, Japan). Samples were visualized in Camscan-S2 (Cambridge, UK) scanning electron microscope(117).

### Phage DNA extraction, sequencing and genome analysis

The genome of isolated phage was extracted utilizing phage DNA extraction kit (Norgen Biotek, Canada, Cat No; 46800) according to manufacturer’s protocol. Supernatant with high concentration of phage was used to isolate the genomic DNA. Phage whole-genome sequencing was performed using the Illumina Miseq platform using DNA prep kit (20060060). The genome sequence was assembled using the Spades genome assembler v4.2 with default selection without any careful flag. The contigs were then subjected to checkV1.0.3 (101) for identification and quality assessment of a viral contigs. Thus, identified contigs with complete genome assessment from checkV was then annotated and classified with PhageScope (https://PhageScope.deepomics.org/).

## DATA AVAILABILITY AND STATISTICAL ANALYSIS

The original results of the study are included in the article. Additional data are included as supplementary data. Further inquiries can be directed to the corresponding author(s). The complete genome sequences of Phage_KP6697_Omshanti have been deposited in GenBank database under the accession number _SRX27372467__and SRA ID number 36925110. Received: 1/11/2025; Accepted: 1/11/2025; Published online: 1/16/2025. Similarly, the complete genome sequences of *Klebsiella pneumoniae* (KP6697) have also been deposited in GenBank database under accession number SRX26555504 and SRA ID number 35877059. Received: 10/16/2024; Accepted: 10/16/2024; Published online: 10/30/2024

## ACKNOWLEDGEMENTS

The authors would like to thank all the investigators and express sincere gratitude to Associate Prof. Dr. Giri Raj Tripathi for his support, Dr. Yuba Raj Pokharel for assistance with the SEM operation at SAIF, AIIMS, Dr. Pramod Poudel and Dr. Pramod Aryal for research guidance, Prajwal Rajbhandari for providing us molecular reagents and the UGC, Nepal for their unwavering support in carrying out this research.

## AUTHOR CONTRIBUTIONS

Dipendra Kumar Mandal: Manuscript conceptualization and funding acquisition, project supervision and data management, methodology and investigation, manuscript writing and editing Gorkha Raj Giri, Roshan Nepal, Gun Raj Dhungana and Pragati Pradhan: Conceptualization, Primer design, Manuscript editing, Method validation, PCR assay Rojina Pandey, Elisha Upadhaya, Puja Dahal, Abdul Rehaman, Sudip Timilsina, Pragya Sapkota, Shobha Amagain, Keshab Gorathoki, Bijita Neupane, Sangharsika Chaudhary, Sushila Thapa, David Pun, Kundan Khadka and Gaurav Adhikari: Data compilation and interpretation, Methodology and investigation Rajindra Napit: Manuscript conceptualization, editing, analysis and visualization, supervision, review, clinical interpretation and writing of results Binod Manandhar: NCBI submission Rajani Malla & Krishana Das Manandhar: supervision for analysis, critical reading, editing, review and improvement of the manuscript. All authors have read and approved the final version of the manuscript.

## Supplemental Materials

The online version contains supplementary material available at ….

## Ethical approval

This study was approved by the Institutional Review Board (IRB) of National Health Research Council (NHRC), Nepal (Ref. No.: 1304). The study does not involve any human and/or animal subjects. The clinical isolates were retrospectively collected from hospital was deidentified and no personally identifiable patient information was disclosed to the researchers. The patient data were not included in this study, therefore informed consent was waived by the NHRC, Nepal.

## Abbreviations

MDR: Multidrug resistance
DLAA: Double layer agar assay
PCR: Polymerase chain reaction
CDS: Coding DNA sequence
tRNA: Transfer RNA
ARG: Antibiotic resistant gene
NCBI: National center for biotechnology information
TEM: Transmission electron microscopy
SEM: Scanning electron microscopy
dsDNA: Double-stranded
DNA G+C: Guanine and cytosine
CRKP: Carbapenem resistance *Klebsiella pneumonia*

---

**Fig S1:** Detection of Metallo-β-lactamase (MBL) production

**Fig S2:** Phage isolation and plaque morphology.

**Table S1:** List of metabolic pathways detected with numbers of associated genes in isolate KP6697 and their respective KO numbers.

**Table S2:** List of social genes found in the *K. pneumoniae* (KP6697) KO identification number and KO definitions.

